# Cooperative multi-view integration with Scalable and Interpretable Model Explainer

**DOI:** 10.1101/2025.01.11.632570

**Authors:** Jerome J. Choi, Noah Cohen Kalafut, Tim Gruenloh, Corinne D. Engelman, Tianyuan Lu, Daifeng Wang

## Abstract

Single-omics approaches often provide a limited perspective on complex biological systems, whereas multi-omics integration enables a more comprehensive understanding by combining diverse data views. However, integrating heterogeneous data types and interpreting complex relationships between biological features—both within and across views—remains a major challenge. To address these challenges, we introduce COSIME (Cooperative Multi-view Integration with a Scalable and Interpretable Model Explainer). COSIME applies the backpropagation of a learnable optimal transport algorithm to deep neural networks, thus enabling the learning of latent features from several views to predict disease phenotypes. It also incorporates Monte Carlo sampling to enable interpretable assessments of both feature importance and pairwise feature interactions for both within and across views. We applied COSIME to both simulated and real-world datasets—including single-cell transcriptomics, spatial transcriptomics, epigenomics and metabolomics—to predict Alzheimer’s disease-related phenotypes. Benchmarking of existing methods demonstrated that COSIME improves prediction accuracy and provides interpretability. For example, it reveals that synergistic interactions between astrocyte and microglia genes associated with Alzheimer’s disease are more likely to localize at the edges of the middle temporal gyrus. Finally, COSIME is also publicly available as an open source tool.

## Main

Single-omics approaches offer limited insight into complex biological systems, as each molecular layer (e.g., genomics, transcriptomics, or metabolomics) functions in coordination with others. Multi-view data integration combines multiple layers to identify novel biomarkers, understand disease mechanisms, and improve phenotype prediction^1,2^. Machine learning can efficiently analyze large-scale multi-view data, modeling complex cross-view interactions that influence phenotypes.

Several methods have been developed for multi-view data integration. Cooperative learning^3^ is a supervised learning method that combines squared-error loss with an agreement penalty encouraging feature similarity across views. The penalty strength is fixed and manually tuned, limiting dynamic adaptation and end-to-end learning of cross-view interactions. DIABLO^4^ identifies shared structure across omic layers through supervised feature selection, but assumes homogeneity in their association with the outcome, which may not hold in heterogeneous datasets. MOFA+^5^ is an unsupervised factor analysis method that models latent linear structures across views, but it lacks mechanisms for interaction analysis or interpretability of prediction features. Furthermore, these models typically assume linear relationships and do not leverage deep learning or optimal transport.

Interpretability is another major challenge. SHAP (SHapley Additive exPlanations)^6^ quantifies feature attribution but is computationally expensive for complex models and can assign interaction contributions only for tree-based models. LIME^7^ provides local interpretability for individual predictions but lacks global explanations and does not account for feature interactions.

To address these limitations, we introduce Cooperative Multi-view Integration with Scalable and Interpretable Model Explainer (COSIME). COSIME consists of two components: (1) an integration component that uses deep neural encoders and Learnable Optimal Transport (LOT) to align views in a joint latent space and predict disease phenotypes; (2) an interpretation component that estimates feature importance (FI) and pairwise interactions using Shapley values and Shapley-Taylor indices. We evaluate COSIME on simulated and real-world datasets to predict Alzheimer’s disease (AD) phenotyes using transcriptomics, metabolomics, epigenomics, and spatial transcriptomics, and we demonstrate its strong predictive performance, flexibility and interpretability.

## Results

### Overview of COSIME

#### COSIME framework

Fig. 1 provides an overview of COSIME. The first component involves integrating multi-view data for disease phenotype prediction by leveraging deep learning-based encoders and learnable optimal transport (LOT), which model both linear and nonlinear relations more flexibly than traditional linear methods. COSIME effectively captures the complex, multi-layered interactions between different omic modalities—such as transcriptomics and epigenomics, transcriptomics and spatial transcriptomics, and transcriptomics and metabolomics—while preserving the distinct characteristics of each data type.

**Fig. 1:**
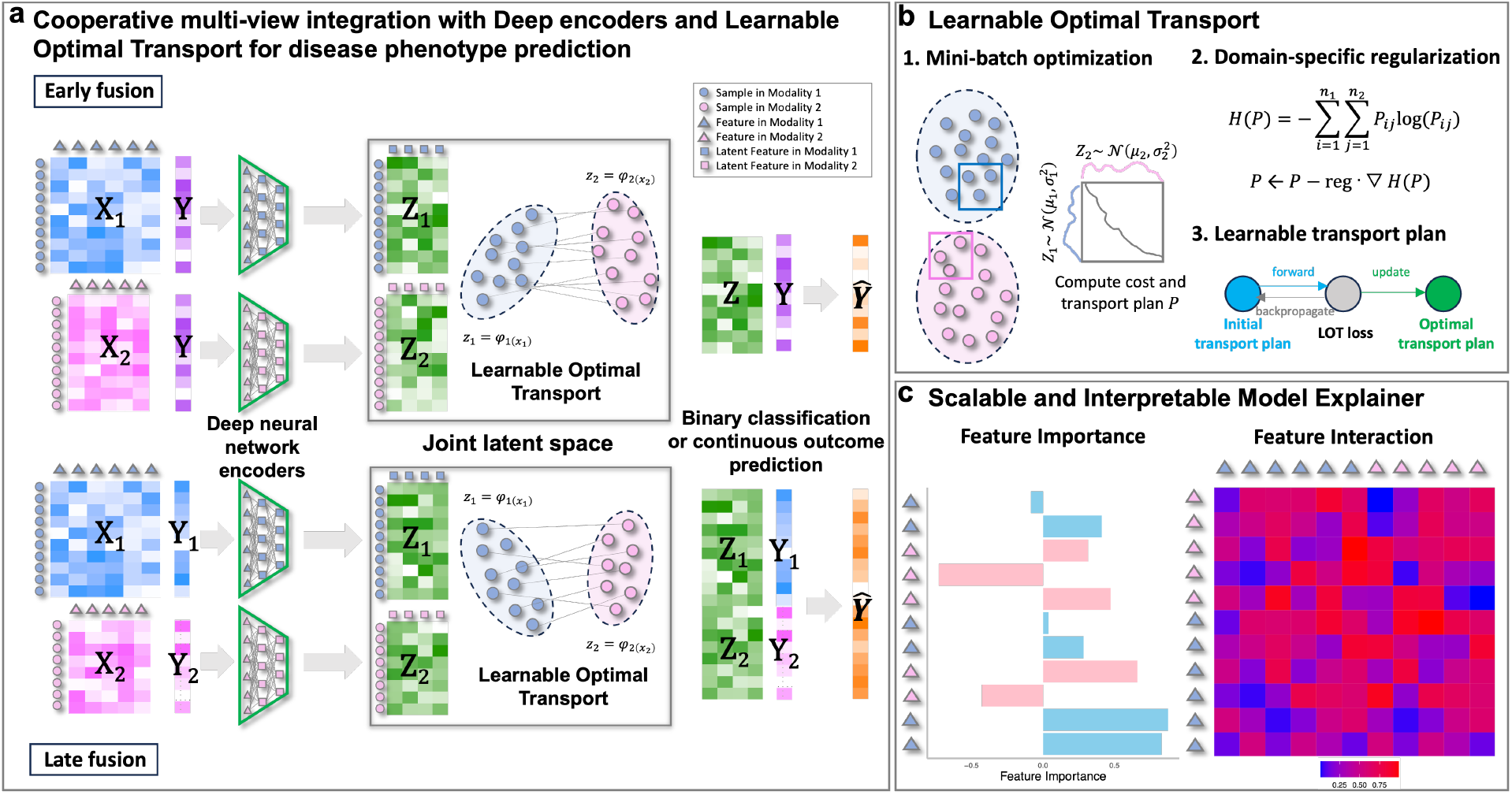
**a**, COSIME integrates multi-view data for disease phenotype prediction through a three-step process: Each data view is passed through separate deep neural network encoders (deep encoders). (2) The learnable optimal transport (LOT) aligns and merges these features into a joint latent space. (3) The integrated latent representation is then used to predict disease phenotypes. **b**, LOT integrates heterogeneous datasets by offering three key advantages: (1) a learnable transport plan that adapts during training, (2) mini-batch processing for scalability, and (3) domain-specific regularization to enhance alignment between source and target distributions. **c**, The scalable and interpretable model explainer interprets model predictions by identifying important features and feature interactions. It quantifies both individual feature importance (FI) and pairwise feature interactions to reveal synergistic or antagonistic effects on the model’s output.

LOT offers three main advantages over conventional optimal transport methods. First, it uses a learnable transport plan, enabling dynamic alignment of heterogeneous data. Second, LOT supports mini-batch processing, making it scalable for large datasets while maintaining computational efficiency. Finally, LOT can incorporate domain-specific regularization, allowing improved handling of view-specific differences and robustness over traditional methods.

The second component of COSIME focuses on computing feature importance (FI) values for each view, as well as pairwise interaction values for both within-view and across-view interactions. To achieve this, COSIME employs the Scalable and Interpretable Model Explainer, which leverages the Shapley values and Shapley-Taylor indices^8^ to compute both feature importance and pairwise feature interactions, respectively. Monte Carlo sampling and batch processing enable scalable computation and enhance interpretability. Additionally, it allows for the identification of the directionality of interactions—whether they exhibit synergism (complementary effects) or antagonism (conflicting effects)—offering deeper insights into how features interact in the model.

COSIME achieves strong predictive performance by modeling across-view relationships within a shared latent space. It integrates heterogeneous data and identifies feature importance and interactions to improve accuracy and interpretability. While these two main components of COSIME can be used together for both enhanced prediction and interpretability, they are also designed to be used independently depending on the specific needs of the analysis. Together, they address the challenges of heterogeneous data and complex feature interactions, enabling biomarker discovery and mechanistic understanding.

In the following sections, the performance metrics were compared across different methods using the mean and ±1.96 times the standard deviation from 5-fold cross-validation. For the best-trained models across the different multi-view datasets, we computed feature importance for each data view and feature interaction values both within and across views. All results for the multi-view predictions using COSIME models and other methods are provided in the **Supplementary Data 1** and **2**. Feature importance and interactions for the simulated data can be found in the **Supplementary Data 3–6**.

### Simulation study

Multi-view data were generated under different signal levels (high and low) and outcome types (binary and continuous). COSIME early fusion performed best on high-signal binary data, outperforming CL^3^, DIABLO^4^, and MOFA+^5^ with logistic regression (AUROC: 0.845 ± 0.026, AUPRC: 0.854 ± 0.029, accuracy: 0.754 ± 0.054). COSIME late fusion also outperformed benchmarks (AUROC: 0.828 ± 0.021, AUPRC: 0.853 ± 0.021, accuracy: 0.761 ± 0.019) (**Fig. 2a**). For low-signal binary data, late fusion yielded the best results (AUROC: 0.737 ± 0.017, AUPRC: 0.736 ± 0.025, accuracy: 0.640 ± 0.052), and early fusion still surpassed all benchmarks.

**Fig. 2:**
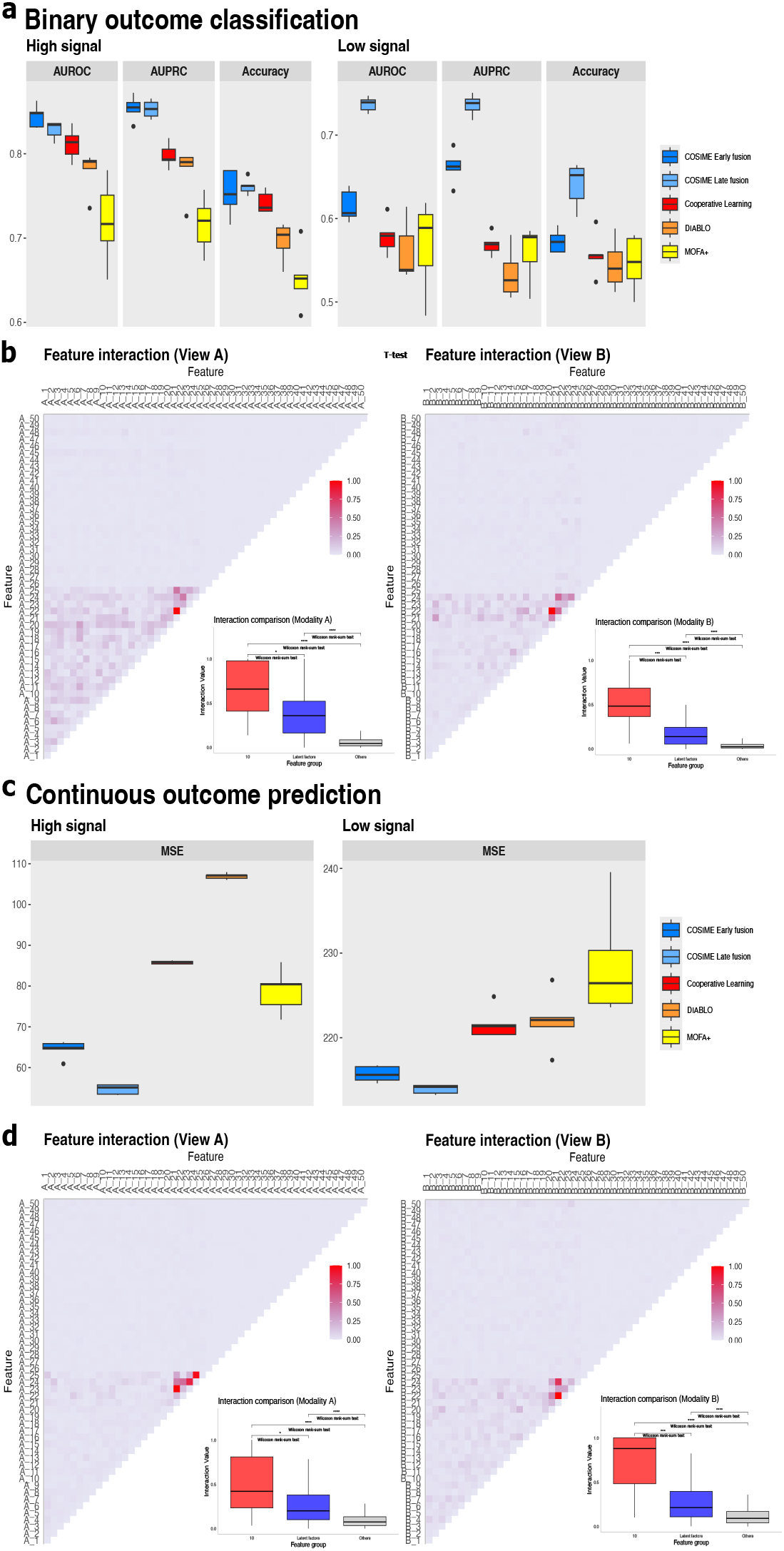
Prediction performance and feature interaction from simulated data. **a**, Predictive performance for binary outcome classification using COSIME early fusion, COSIME late fusion, CL, DIABLO, and MOFA+ with logistic regression on both high-signal and low-signal multiview datasets. **b**, Heatmaps depicting the pairwise interactions of the first 50 features from views A and B for binary outcome prediction, along with box plots and two-sided Wilcoxon rank-sum tests for three interaction groups: (Group 1) 10 interaction terms artificially introduced during data generation, (Group 2) 1,790 interactions involving latent features, and (Group 3) 3,150 other interactions. The box plots display the distribution of interaction values, including the first quartile, median, and third quartile. Asterisks indicate statistical significance: * for *p*–value < 0.05 and **** for *p*–value < 0.0001. **c**, Prediction performance for continuous outcome prediction using COSIME early fusion, COSIME late fusion, CL, DIABLO, and MOFA+ with regression on both high-signal and low-signal multi-view datasets. **d**, Heatmaps depicting the pairwise interactions of the first 50 features from views A and B for continuous outcome prediction, along with box plots and pairwise two-sided t-tests for three interaction groups: (Group 1) 10 interaction terms artificially introduced during data generation, (Group 2) 1,790 interactions involving latent features, and (Group 3) 3,150 other interactions. The box plots display the distribution of interaction values, including the first quartile, median, and third quartile. Asterisks indicate statistical significance: *** *p*–value < 0.001 and **** for *p*–value < 0.0001.

Feature interaction values from COSIME early fusion (high-signal binary data) were grouped as follows: (1) artificially introduced, (2) involving latent features and (3) others. Scaled interactions for the first 50 features are shown in **Fig. 2b**. Wilcoxon rank-sum tests show Group 1 > Group 2 (*p <* 0.05), Group 1 > Group 3 (*p <* 0.0001), and Group 2 > Group 3 (*p <* 0.0001), in both views A and B. For high-signal continuous outcomes, COSIME achieved the lowest MSE (54.701 ± 2.391), outperforming CL, DIABLO, and MOFA+ (MSE: 64.490 ± 4.130) (**Fig. 2c**). On low-signal data, COSIME early fusion again performed best (MSE: 215.707 ± 1.833), while late fusion outperformed benchmarks (MSE: 218.425 ± 1.099). Interaction analysis for high-signal continuous data (top 50 features in **Fig. 2d**) shows consistent results: Group 1 > Group 2 (*p <* 0.05) and Group 3 (*p <* 0.0001); Group 2 > Group 3 (*p <* 0.0001) across both views. Overall, COSIME consistently outperforms all benchmarking methods and recovers synthetic interactions introduced during data generation.

To confirm the superiority of COSIME for multi-view integration and outcome prediction, we compared its performance against other methods. The results are presented in **Supplementary Table 1** and **Supplementary Figs. 2 and 3**. COSIME with both early and late fusion outperforms its single-view methods, single-view regressions and benchmark methods, including aligned OT with regression and single-cell multi-omic integration with optimal transport (SCOT) with regression. These findings highlight the strength of the COSIME integration and show that LOT in COSIME surpasses other OT-based approaches.

To illustrate the computational efficiency of COSIME, we report runtimes for various dataset sizes in **Supplementary Table 3**.

To justify the superiority of COSIME for multi-view integration and outcome prediction, we compared its performance against other methods. Results are presented in the **Supplementary Table 1** and **Supplementary Figs. 2** and **3**. COSIME with both early and late fusion outperformed its single-view methods, single-view regressions, and benchmark methods including Aligned OT with regression and SCOT OT with regression. These findings highlight the strength of COSIME’s integration and show that LOT in COSIME surpasses other optimal transport-based approaches.

To provide guidance on the computational efficiency of COSIME, we report run times for varying dataset sizes in the **Supplementary Table 3**.

### Cognitive diagnosis from transcriptomic and metabolomic data

Multi-view data from scRNA-seq of astrocytes and metabolomics were used to test the prediction performance of COSIME models. COSIME early fusion with matched samples (n=2,286) achieved the best performance for binary classification of Alzheimer’s disease (AD) cognitive diagnosis using multi-view datasets (astrocytes and metabolomics) (AUROC: 0.792 ± 0.021, AUPRC: 0.810 ± 0.022, accuracy: 0.732 ± 0.025). To leverage all available samples (4292 metacells in transcriptomics and 2286 samples in metabolomics) across views, we also applied COSIME late fusion for unmatched samples, which yielded comparable results (AUROC: 0.807 ± 0.013, AUPRC: 0.832 ± 0.016, accuracy: 0.723 ± 0.027) (**Fig. 3a**).

**Fig. 3.**
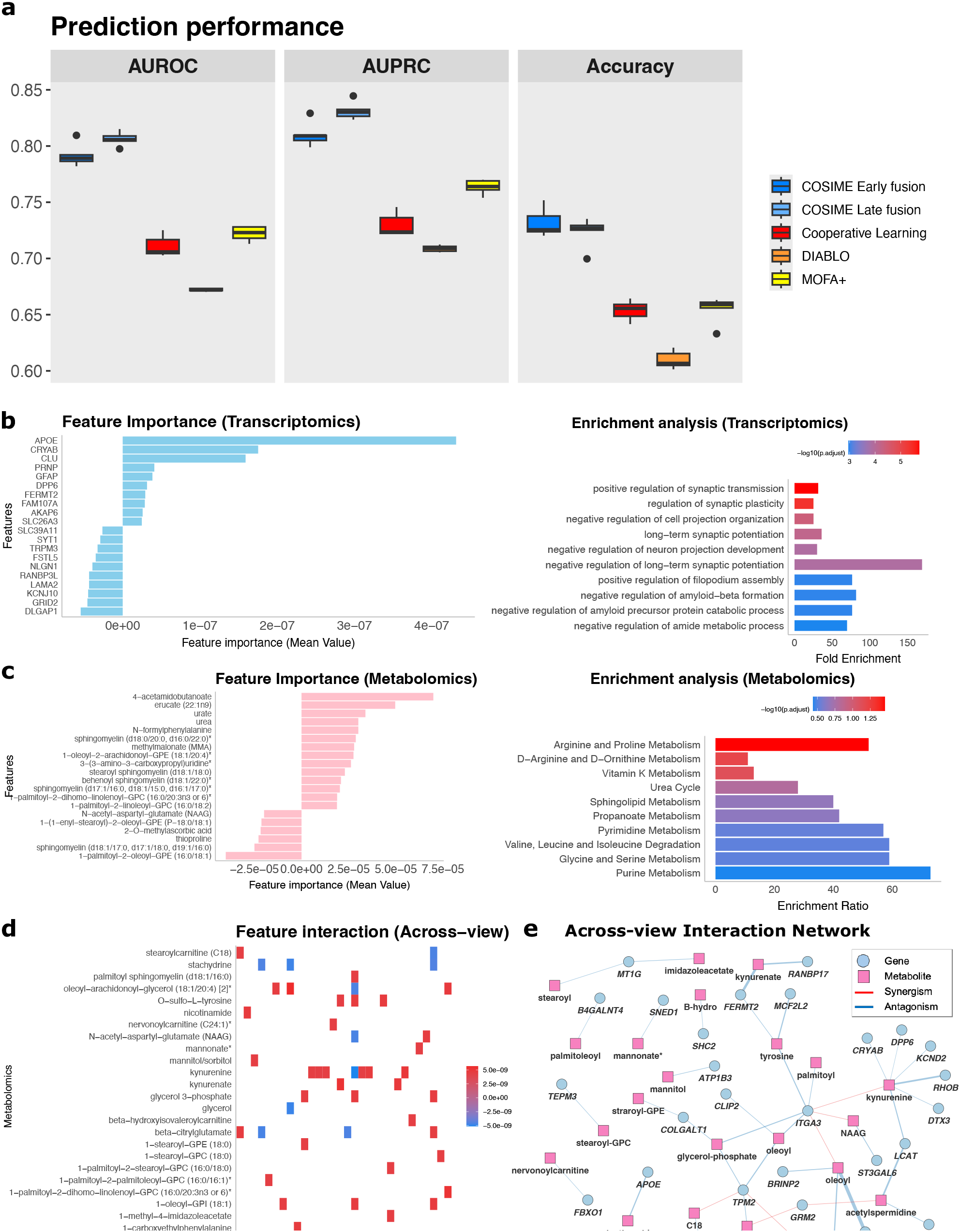
Classifying cognitive diagnoses for AD from transcriptomic (astrocytes) and metabolomic data. **a**, Predictive performance using COSIME early fusion, COSIME late fusion, CL, DIABLO, and MOFA+ with logistic regression. **b**, Bar plots showing the top 20 prioritized genes (absolute feature importance values) and enrichment analysis for those genes. **c**, Bar plots showing the top 20 prioritized metabolites (absolute feature importance values) and enrichment analysis for those metabolites. **d**, Heatmap showing the top 50 pairwise across-view (gene-metabolite) interactions. **e**, Across-view interaction network showing relationships between the top 50 pairwise across-view (gene-metabolite) interactions.

We also compared its performance against other methods. Overall, COSIME with both early and late fusion outperformed its single-view methods, single-view regressions, and existing benchmark methods including Aligned OT with regression and SCOT OT with regression. Results are presented in the **Supplementary Table 2** and **Supplementary Fig. 4**.

FI values were used to prioritize features. The top 20 most important genes (based on absolute values) from the transcriptomic data are displayed in **Fig. 3b**, and enrichment analysis reveals that these genes are strongly associated with AD. Notably, processes such as synaptic transmission, amyloid-beta (*Aβ*) formation, and synaptic plasticity are strongly implicated in AD. They represent crucial aspects of AD pathophysiology^9,10^.

**Fig. 3c** shows the top 20 most important metabolites (ranked by absolute feature importance values) from the metabolomic data. Enrichment analysis of these metabolites highlights metabolic pathways such as arginine and proline metabolism vitamin K metabolism, and sphingolipid metabolism. These pathways may contribute to AD by exacerbating the brain’s energy deficit, promoting oxidative damage, and driving neuroinflammation^11^.

Pairwise feature interaction values are computed, and the top 50 across-view feature interaction values are shown in **Fig. 3d**. *APOE* has synergistic effects with nicotinamide. The *APOE* ε4 allele is the strongest genetic risk factor for late-onset AD, promoting amyloid-beta aggregation, neuroinflammation, and synaptic dysfunction^12^. Nicotinamide, a form of vitamin B3 and a precursor to NAD^+^, plays a critical role in mitochondrial function, oxidative stress response, and DNA repair. Altered nicotinamide metabolism in AD reflects disrupted energy homeostasis and neurodegenerative stress. Together, elevated *APOE* expression and dysregulated nicotinamide metabolism may synergistically exacerbate AD pathology. *APOE*driven neuroinflammation and amyloid deposition may increase cellular demand for NAD^+^, while impaired nicotinamide pathways may fail to meet this demand, leading to mitochondrial dysfunction and neuronal vulnerability. Their combined dysregulation amplifies the biological signals of neurodegeneration, contributing to a stronger predictive signature for AD in machine learning models.

*ATP5PF* shows antagonistic effects with stachydrine (**Fig. 3d**). *ATP5PF* is a subunit of mitochondrial ATP synthase essential for cellular energy production through oxidative phosphorylation. In AD, reduced *ATP5PF* expression reflects mitochondrial dysfunction and impaired energy metabolism, both of which are key features of neurodegeneration^13^. Stachydrine is a plant-derived metabolite known for its antioxidant and mitochondrial protective effects. It has been shown to reduce oxidative stress, preserve mitochondrial function, and support cellular energy homeostasis in various disease models^14^. They may exhibit an antagonistic relationship in AD, where high levels of stachydrine may counteract the detrimental effects of *ATP5PF* downregulation. This suggests that stachydrine acts as a compensatory factor against mitochondrial dysfunction, potentially buffering the impact of energy failure in the brain.

Furthermore, *TPM2* and *ITGA3* may serve as important hub genes in the across-view network (**Fig. 3e**). These two genes are likely involved in astrocyte function under both normal and pathological conditions, particularly through their roles in inflammation, actin remodeling, and neurovascular interactions. Kynurenine may be a hub metabolite, interacting with genes. Through its role in neuroinflammation and oxidative stress, kynurenine may antagonize the function of key astrocyte genes that support neuronal survival, metabolic stability, and synaptic homeostasis. In AD, elevated kynurenine may disrupt astrocyte-mediated neuroprotection, leading to an inverse or antagonistic relationship with homeostatic astrocyte genes^15^.

Feature importance and interactions for the ROSMAP binary classification early fusion are in the **Supplementary Data 7**.

### Cognitive status from transcriptomic and epigenomic data

Multi-view data from both scRNA-seq and scATAC-seq of oligodendrocytes were utilized to evaluate the predictive performance of COSIME models. COSIME early fusion performed the best for binary classification of cognitive status (dementia) using multi-view datasets (oligodendrocytes) (AUROC: 0.874 ± 0.011, AUPRC: 0.805 ± 0.013, accuracy: 0.794 ± 0.041) (**Fig. 4a**).

**Fig. 4.**
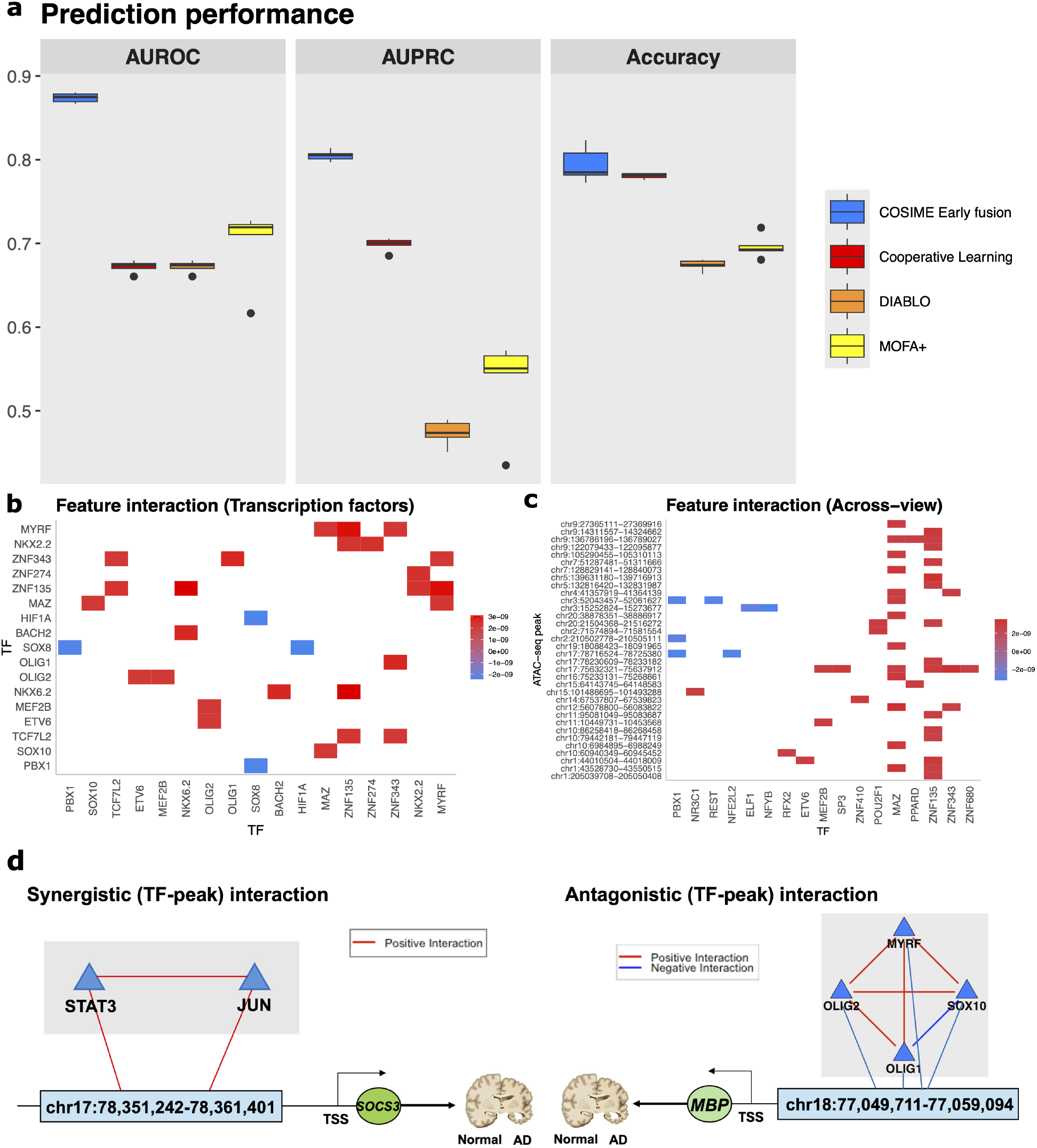
Classifying dementia from transcriptomic (oligodendrocyte) and epigenomic (oligodendrocyte) data. **a**, Predictive performance using COSIME early fusion, CL, DIABLO, and MOFA+ with logistic regression. **b**, Heatmap showing the top 50 pairwise TF interactions. **c**, Heatmap showing the top 50 pairwise across-view (TF-ATAC peak) interactions. **d**, Diagrams illustrating examples of synergistic and antagonistic relationships between a TF-ATAC peak regulating a target gene (TG) at their transcription start site (TSS) and the AD phenotype.

We investigated pairwise interactions between the ten known key TFs involved in oligodendrocytes^16^. For instance, synergistic interactions were identified between SOX10 and MAZ as well as between MYRF and MAZ. Specifically, SOX10 is critical for oligodendrocyte differentiation^17^. MAZ is involved in regulating genes related to cell proliferation, differentiation, and survival^18^. Their co-expression may suggest a compensatory mechanism for myelin repair in dementia. MYRF is another key player in myelination^19^. Its synergism with MAZ may similarly support remyelination in disease contexts (**Fig. 4b**).

Moreover, antagonistic interaction is identified between SOX8 and HIF1A. SOX8 regulates development and differentiation^20^, while HIF1A is a stress-related factor^21^ that could suppress differentiation and thereby contribute to dementia pathology. SOX8 also shows antagonism with PBX1, a TF involved in neural cell development^22,23^. This interaction may reflect a regulatory imbalance affecting myelin integrity (**Fig. 4b**).

Based on the top fifty across-view absolute interaction values, we identified several TFs interacting with multiple oligodendrocyte-specific regulatory regions. MAZ showed synergistic interactions with ATAC peak regions mapping to genes like *PTPRF*^24^, *ERBB3*^25^, *GRID1*^26^, *LIMCH1*^27^, *FLNC*^28^, and *DUSP7*^29^. These interactions suggest MAZ may influence gene networks relevant to neurodegeneration. ZNF135 also showed synergism with peaks linked to *NKX2-2*^30^, *PTPRF*^24^, *CNTN2*^31^, *PSD2*^32^, and *SLC6A9*^33^, associated with neural development and synaptic function (**Fig. 4c**).

In contrast, PBX1 antagonized open chromatin regions and may reduce accessibility, impairing activation of protective genes such as *CPS1*^34^ and *CYTH1*^35^. This may reflect loss of protective oligodendrocyte mechanisms (**Fig. 4c**).

STAT3 and JUN exhibit synergistic activity in the chr17:78,351,242-78,361,401 region containing *SOCS3*. STAT3 regulates JAK-STAT signaling, immune response, and differentiation^36,37^, while JUN regulates stress and inflammatory gene expression^38,39^. Both TFs are known to regulate *SOCS3*, a gene upregulated in dementia^40,41^. Their coordination may contribute to *SOCS3* dysregulation and downstream inflammation (**Fig. 4d**).

The four key TFs — SOX10^17^, MYRF^17^, OLIG1^42^, and OLIG2^42^ — show synergistic relationships, except SOX10 and OLIG1, which are antagonistic. These TFs regulate oligodendrocyte differentiation and myelination. Their synergy may contribute to oligodendrocyte dysfunction and myelin damage in dementia. All four TFs antagonistically interact with an ATAC peak near *MBP* (chr18:77,049,711-77,059,094), a key myelin gene^43^, suggesting suppressed *MBP* expression (**Fig. 4d**).

Feature importance and interactions for the SEA-AD scRNA-seq and scATAC-seq (oligodendrocytes) early fusion are included in the **Supplementary Data 8**.

### Predicting AD progression scores from transcriptomic data

To assess the predictive capability of COSIME models, multi-view data combining scRNA-seq of microglia and single-cell spatial transcriptomics (MERFISH) of astrocytes were applied. COSIME early fusion performed better than the other benchmarking methods for continuous outcome prediction of Alzheimer’s disease progression scores using multi-view datasets (microglia and astrocytes) (MSE:0.033 ± 0.005) (**Fig. 5a**).

**Fig. 5.**
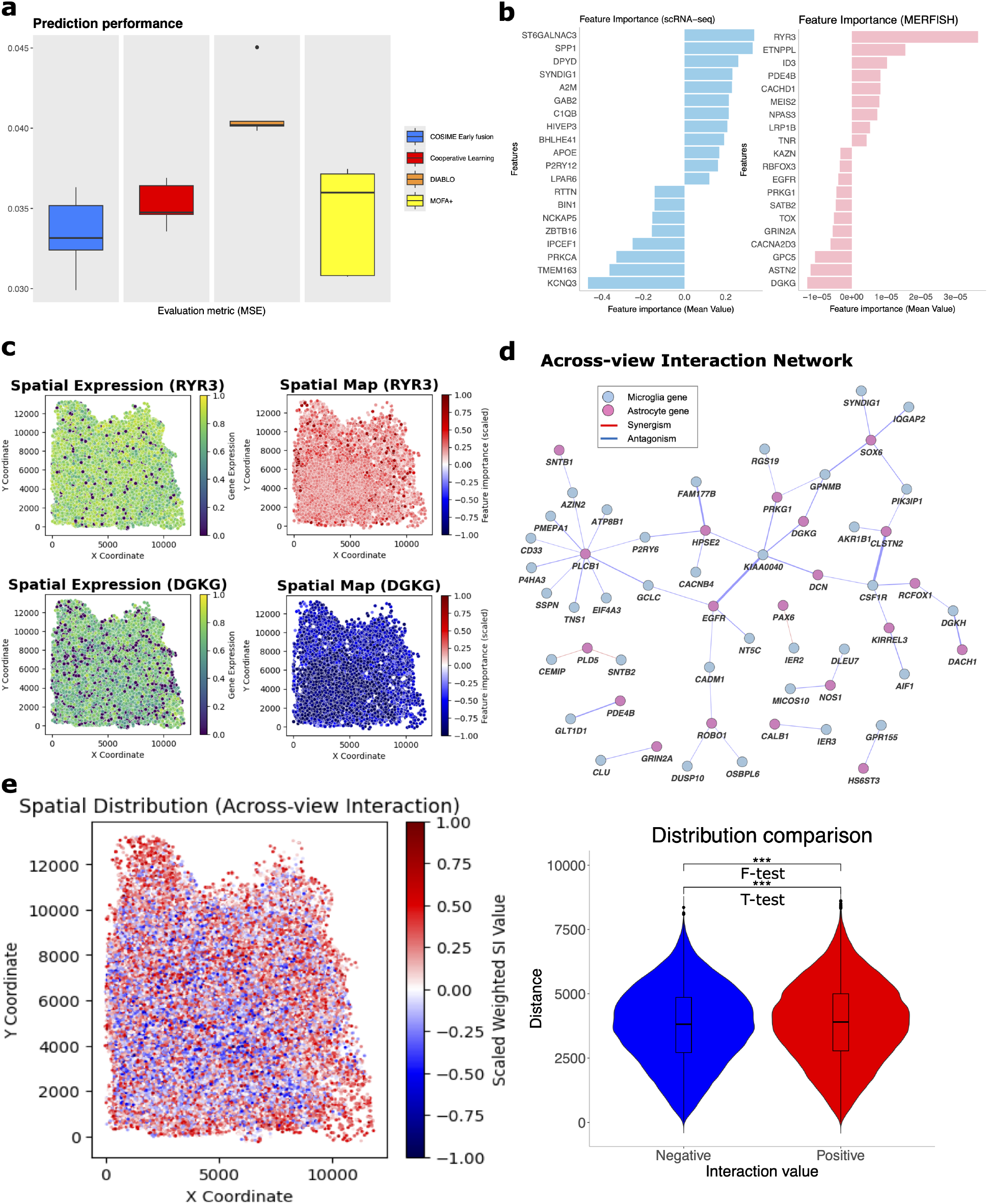
Predicting AD progression scores from transcriptomic (microglia) and spatial transcriptomic MERFISH (astrocytes) data. **a**, Predictive performance using COSIME early fusion, CL, DIABLO, and MOFA+ with regression. **b**, Bar plots showing the top 20 prioritized genes (absolute feature importance values) in each view. **c**, Spatial plots showing aexpression level and importance values across cells for *RYR3* and *DGKG*. **d**, Across-view interaction network showing relationships between the top 50 pairwise across-view (gene-gene) interactions. **e**, Spatial distribution of all across-view interaction values colored by the directionality and the distribution of 80,072 positive and 68,619 negative interaction values, with statistical tests (two-tailed t-test and two-tailed F-test) applied to assess differences between two groups. The violin plots including box plots display the distribution of interaction values in the two groups. Asterisk indicate statistical significance: *** for *p*–value < 0.001.

In microglia, several genes with positive feature importance are linked to AD through roles in neuroinflammation, clearance, and activation. *SPP1* is a microglial marker involved in inflammatory response and amyloid plaque phagocytosis^44^. *APOE*, particularly the E4 allele, is a major AD risk gene affecting clearance and neuroinflammation^12,45^. *C1QB*, part of the complement system, tags damaged neurons for microglial clearance^46,47^. Genes with negative feature importance like *BIN1*, involved in synaptic regulation and endocytosis^48^, may be downregulated in AD microglia. **Fig. 5b** also shows that in astrocytes, genes with negative importance like *RYR3* may reflect impaired synaptic regulation via calcium signaling^49,50^. *DGKG*, with positive importance, is involved in lipid metabolism and inflammation^51^. Its dysregulation may affect glutamate signaling, contributing to synaptic damage. Top enriched pathways (**Supplementary Fig. 1**) are largely AD-relevant. We examined spatial distribution of cells based on *RYR3* expression and importance values (**Fig. 5c**). Cells with top 25% *RYR3* feature importance values are significantly central (t-test). Similarly, negatively important *DGKG* cells are also more central (t-test). Together, this suggests central enrichment of downregulated genes and peripheral activity of upregulated genes (**Fig. 5c**). We constructed a gene-gene interaction network from the top 50 across-view interaction values. Astrocyte genes *PLCB1* and *HPSE2* are key hubs linking multiple microglial genes. *PLCB1* regulates phosphatidylinositol signaling and inflammation^52^. *HPSE2* modulates extracellular matrix remodeling and immune responses^53^ (**Fig. 5d**). A spatial map based on across-view interactions was generated (**Fig. 5e**). Both t-test and F-test reject the null (t-stat: 18.783, F-stat: 1.242,), indicating distinct spatial distributions. Synergistic interactions are more peripheral, while antagonistic ones are central (**Fig. 5e**). In AD, the middle temporal gyrus (MTG) is vulnerable to early pathological changes including amyloid and tau accumulation^54,55^. Peripheral MTG regions may experience stronger glial activation and synaptic stress^56,57^, making them potential early hotspots for AD-related gene activity. While microglia and astrocytes are both essential for maintaining brain function, their interactions at the gene level can be either synergistic or antagonistic^58–60^.

FI and interactions for the SEA-AD scRNA-seq (microglia) and MERFISH (astrocytes) early fusion are shown in the **Supplementary Data 9**.

## Discussion

We introduce COSIME (Cooperative Multiview Integration with Scalable Interpretable Model Explainer) and apply it to various multi-view datasets, including simulated data, transcriptomics-metabolomics, transcriptomics-epigenomics, and transcriptomics-spatial transcriptomics. COSIME has two key components: first, it integrates multi-view data for disease phenotype prediction by leveraging deep encoders and LOT techniques, combining both unsupervised and supervised learning. Second, it computes feature importance for each view and pairwise interactions for both within- and across-view features.

COSIME addresses several challenges in the existing literature. First, it handles both linear and nonlinear relationships and captures feature interactions. The deep encoders generate embeddings without relying on linear assumptions, which can then be used with linear or nonlinear models. Second, COSIME supports matched and unmatched samples via separate deep encoders, applying early or late fusion accordingly. LOT is a specialized technique that aligns embeddings into a joint latent space. Third, LOT is a unique technique that aligns latent distributions using a learnable transport plan dynamically optimized during training. This allows flexible adaptation to data structure and improves downstream prediction tasks. Fourth, COSIME efficiently computes FI and feature interactions via Monte Carlo sampling and batch processing. Lastly, COSIME identifies the directionality of interactions—synergism or antagonism—enhancing interpretability and providing intuitive output matrices.

COSIME incorporates LOT, which is developed for integrating two heterogeneous multiview datasets in predictive modeling. This capability is particularly important given the nonlinear nature of omics data—such as transcriptomics, metabolomics, epigenomics, and spatial transcriptomics. Unlike existing methods such as Cooperative Learning^3^, DIABLO^4^, and MOFA+^5^, which are not specifically designed to capture nonlinear relations, COSIME (both early and late fusion) effectively captures nonlinear dependencies and consistently outperforms these methods in phenotype prediction. In addition, COSIME outputs both FI and interaction values from any complex models, whereas SHAP^6^ provides interaction matrices only for tree-based models and LIME^7^ is limited to computing feature attributions only from simpler models. Beyond these analytical capabilities, COSIME is user-friendly, allowing users to train a model, compute feature relevance with a pretrained model, or combine both, while supporting flexibility in fusion strategy and output type.

We show COSIME outperforms well-known multi-omics models in integration and prediction, while also providing interpretable insights. However, there is room for further development. First, at present, COSIME computes only pairwise feature interactions; higher-order interactions may yield richer explanations but would require more advanced approaches. Second, selecting the number of hidden layers and the size of the latent space is not trivial. Although the latent dimensionality was fine-tuned, optimizing the network architecture will be explored in future work. Third, while our framework currently supports two data views, it can be extended to more views by adding separate encoders and aligning them in a shared latent space. For example, integrating transcriptomics, chromatin accessibility, and DNA methylation^61^ would involve applying LOT pairwise (e.g., transcriptomics–epigenomics, epigenomics–methylation, and transcriptomics–methylation) and jointly optimizing their alignment. This also enables integration of modalities such as Hi-C^62^ with suitable representations. Lastly, COSIME could also be applied to other brain-related diseases, such as neuropsychiatric disorders^63,64^, when largescale multi-omics datasets become available. It can also integrate with methods for multi-view imputation^65–67^, enhancing its capacity to handle missing data and broadening its applicability.

## Methods

### COSIME overview

Cooperative Multiview Integration with Scalable and Interpretable Model Explainer (COSIME) is a machine learning model that integrates multi-view data for disease phenotype prediction and computes feature importance and interaction scores. By leveraging deep learning-based encoders, COSIME effectively captures the complex, multi-layered interactions between different omic modalities while preserving the unique characteristics of each data type. The integration of LOT techniques aligns and merges heterogeneous datasets, improving the accuracy of modeling across-view relationships in the joint latent space. In addition, COSIME leverages the Shapley-Taylor Interaction Index^8^ to compute feature importance and interaction values, allowing for a deeper understanding of how individual features and their interactions contribute to the predictions (**Fig. 1**).

### Simulated Data

We generated synthetic data by creating two distinct datasets, **X**_1_ and **X**_2_, each with a specified number of features, to simulate a scenario for multi-view analysis. Latent factors, which are unobserved variables that affect the data, were introduced with specified standard deviations (std, std_1_, and std_2_) and were either correlated or uncorrelated depending on the experimental design. The strength of the influence of these latent factors on the multi-view datasets was controlled by the scaling factors scale_1_ and scale_2_. Additionally, interaction effects between the latent factors of **X**_1_ and **X**_2_ were optionally included to introduce non-linear relationships, with the weight of these interactions controlled by the interaction weight parameter.

### Latent Factors and Dataset Generation

We generate the latent factors for the datasets **X**_1_ and **X**_2_ based on the correlation setting. If the latent factors are correlated, we use a shared latent factor matrix **U** while if the latent factors are uncorrelated, we use separate latent factor matrices **U**_1_ and **U**_2_.

#### Latent Factor Generation

The latent factors for both **X**_1_ and **X**_2_ are drawn from normal distributions:

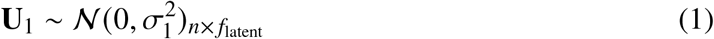

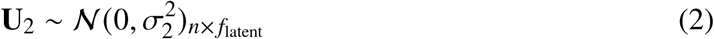

Where:

- *n* is the number of samples.
- *f*_latent_ is the number of latent factors.
- 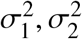 are the variances of the latent factors affecting **X**_1_ and **X**_2_, respectively.

#### Correlated Latent Factors

If the latent factors in the two data views are correlated, we use a shared latent factor matrix **U**, and both datasets **X**_1_ and **X**_2_ are generated as:

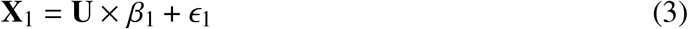

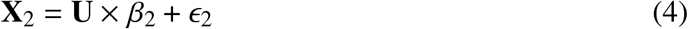

Where:

- **U** is the shared latent factor matrix.
- *β*_1_, *β*_2_ are the weights for **X**_1_ and **X**_2_, respectively.
- *ϵ*_1_, *ϵ*_2_ represent the noise for **X**_1_ and **X**_2_, respectively, and follow:

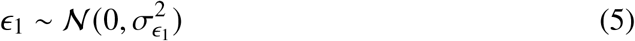

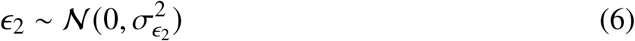

#### Uncorrelated Latent Factors

If the latent factors in the two data views are uncorrelated, we generate separate latent factor matrices **U**_1_ and **U**_2_ for **X**_1_ and **X**_2_, and the datasets are generated as:

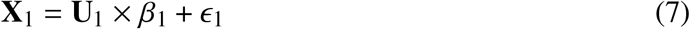

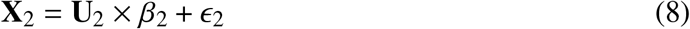

Where:

- **U**_1_ and **U**_2_ are the uncorrelated latent factor matrices for **X**_1_ and **X**_2_.
- *β*_1_, *β*_2_ are the weights for **X**_1_ and **X**_2_, respectively.
- *ϵ*_1_, *ϵ*_2_ represent the noise for **X**_1_ and **X**_2_, respectively, and follow:

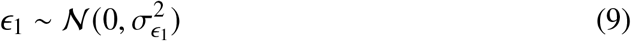

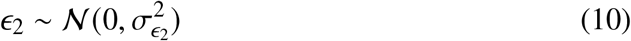

### Interaction Effects

#### Correlated Latent Factors

When the latent factors in the two data views are correlated, the interaction term is generated by considering pairwise combinations of the latent factors. Specifically, the interaction term **z**_interaction_ is calculated as:

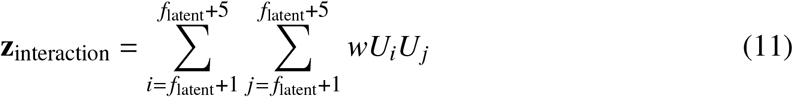

Where:

- *U*_*i*_ and *U* _*j*_ are the elements of the latent factor matrix **U**.
- The interaction term involves the product of latent factors *U*_*i*_ and *U* _*j*_ for each pair.
- The sum is performed over the indices *i* and *j*, which range from *f*_latent_ + 1 to *f*_latent_ + 5, where the interaction is modeled between latent factors.
- *w* is the interaction weight, a constant that determines the strength of the interaction between latent factors.

The interaction term **z**_interaction_, which is given by equation 11, is then added to the feature matrices **X**_1_ and **X**_2_ as follows:

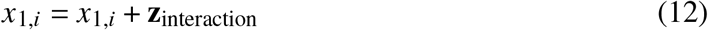

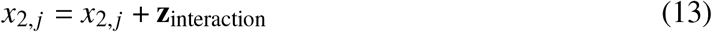

Where equations 12 and 13 show how the interaction term **z**_interaction_ is added to the features *x*_1_ and *x*_2_ in the correlated case.

### Outcome Variable

The outcome variables **y**_1_ and **y**_2_ are generated based on the feature matrices **X**_1_ and **X**_2_, taking into account whether interaction terms are included or not.

#### Correlated Latent Factors

For the case where the latent factors are correlated (i.e., a shared latent factor matrix **U** is used for both **X**_1_ and **X**_2_), the outcome generation is as follows:

- **With Interaction:** When interaction terms are included, the expected outcome *μ*_all_ is computed as the linear combination of the latent factors **U** and the latent factor weights *β*, along with the pairwise interaction terms between the additional latent factors. The interaction term **z**_interaction_, given by equation 11 is used, which is defined as:

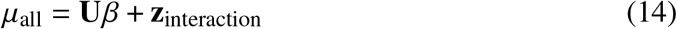
- **Without Interaction:** When no interaction terms are included, the expected outcome is simply the linear combination of the latent factors **U** weighted by *β*:

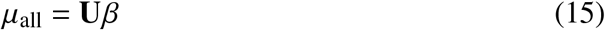

#### Uncorrelated Latent Factors

For the case where the latent factors are uncorrelated (i.e., separate latent factor matrices **U**_1_ and **U**_2_ are used for **X**_1_ and **X**_2_, respectively), the outcome generation is as follows:

- The expected outcome is the sum of the linear combinations of **U**_1_ and **U**_2_ with their respective weights 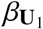and 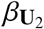:

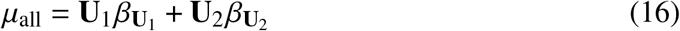

#### Outcome Variable Generation

The outcome variables **y**_1_ and **y**_2_ are generated using the expected outcomes *μ*_all_ computed above, along with Gaussian noise for continuous outcomes or using a logistic function for binary outcomes.

- **Continuous Outcomes:** For continuous outcomes, the outcome variables **y**_1_ and **y**_2_ are generated by adding Gaussian noise to the expected values:

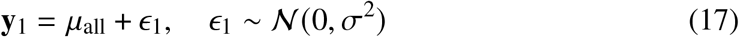

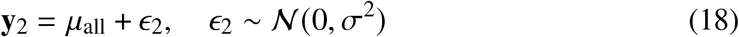

Where *ϵ*_1_ and *ϵ*_2_ are Gaussian noise terms with mean 0 and variance *σ*^2^.

- **Binary Outcomes:** For binary outcomes, the predicted probabilities ŷ_1_ and ŷ_2_ are computed using the logistic function:

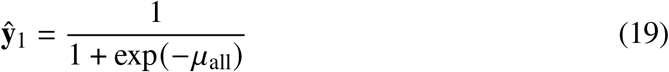

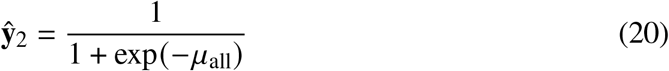

The outcome variables are then generated based on Bernoulli trials with added Gaussian noise to the predicted probabilities:

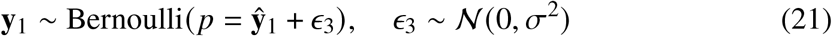

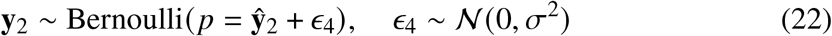

Where *ϵ*_3_ and *ϵ*_4_ are Gaussian noise terms with mean 0 and variance *σ*^2^.

### Experimental Conditions

We generated a total of eight experimental conditions, divided between binary and continuous outcome variables. Each condition included two distinct datasets to facilitate multi-view analysis. The conditions were based on two factors: strength of signal in the data (high vs. low) and the fusion method (early vs. late). For each dataset, we used different random seeds to introduce variability while keeping all other parameters consistent, particularly the latent factor structure **U**. In all datasets, 10 interaction terms were introduced within each view to capture the dependencies between selected features. These interactions model the relationships between certain features within the same view.

The experimental conditions were as follows:

- **Signal Strength and Fusion Type**

- **High Signal (Early Fusion)**: Data were generated with a strong latent factor influence, including interaction effects between features. The latent factor structure **U** and interaction terms remained consistent across both datasets.
- **High Signal (Late Fusion)**: The same strong latent factor influence was used as in the early fusion condition. However, the datasets were generated with different samples, while maintaining consistency in latent factor structure **U** and interaction effects.
- **Low Signal (Early Fusion)**: Data were generated with a weaker latent factor influence. Interaction effects were included, and the latent factor structure **U** and interaction terms remained consistent across both datasets.
- **Low Signal (Late Fusion)**: Data were generated with weak latent factor influence. The two datasets were generated with different samples, while maintaining consistency in latent factor structure **U** and interaction effects.

- **Outcome Type**
- **Binary Outcomes**: Interaction effects were included, and the latent factor structure **U** and interaction terms remained consistent across datasets, based on the signal strength and fusion type.
- **Continuous Outcomes**: Similar to binary outcomes, but the outcome variables were generated using Gaussian noise as detailed in the main text.

### Parameters used to generate the datasets

The following are the parameters used to generate the datasets, based on the notations introduced in the previous sections.

- Sample size: *n* = 1000
- Number of features: 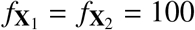
- Number of latent factors: *f*_latent_ = 25 (Including five for interacting features)
- Scaling factors: scale_1_ = scale_2_ = 5 for binary high signal and continuous high signal; scale_1_ = scale_2_ = 10 for binary low signal and continuous low signal
- Latent factor standard deviations: std = 1 (shared standard deviation for latent factors affecting both views if correlated), std_1_ = 1 (standard deviation for latent factors influencing **X**_1_ if uncorrelated), std_2_ = 1 (standard deviation for latent factors influencing **X**_2_ if uncorrelated)
- Factor strength: latent strength = 2 for high signal, latent strength = 1 for low signal
- Noise standard deviation: *σ* = 5 for strong signal conditions, *σ* = 10 for weak signal conditions (binary outcomes), *σ* = 15 for weak signal conditions (continuous outcomes)
- Correlation between two datasets: TRUE if **X**_1_ and **X**_2_ are correlated, FALSE if they are uncorrelated
- Data type: Binary or continuous outcomes for **y**_1_ and **y**_2_
- Interaction effects: TRUE if interactions between features of **X**_1_ and **X**_2_ are included, with interaction weight *w* = 10

- Interaction effects between the 21st and 25th features within each view (**X**_1_ and **X**_2_) were artificially introduced.

Where:

- **X**_1_ and **X**_2_ represent the feature matrices for the two datasets.
- **U** is the shared latent factor matrix when the latent factors are correlated, and **U**_1_ and **U**_2_ are the separate latent factor matrices for the uncorrelated case.
- *β*, 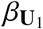, and 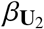are the latent factor weights used to generate the expected outcome variables.
- latent strength indicates the strength of the latent factors’ influence on the outcome variables, with higher values corresponding to stronger signal conditions.
- *σ* denotes the standard deviation of the noise added to the outcome variables.
- *w* is the interaction weight, a constant that controls the influence of interaction terms between features from **X**_1_ and **X**_2_, particularly for the 21st to 25th features in each view.

### Additional datasets

To compare the computing time across datasets of different sizes, we additionally generated the following datasets.

- 5000 samples with 100 features in each view
- 5000 samples with 250 features in each view
- 10000 samples with 500 features in each view

We followed the same settings but proportionally increased the number of latent factors based on the number of features in each scenario.

## Real Data

### Transcriptomics-Metabolomics (ROSMAP)

The *Religious Orders Study* and *Rush Memory and Aging Project* (ROSMAP) are ongoing longitudinal clinical-pathologic cohort studies of aging and AD^68^. The ROSMAP data include a wealth of clinical, cognitive, neuroimaging, genetic, and neuropathological information from older adults, providing invaluable insights into the early stages of AD, cognitive decline, and related neurodegenerative processes.

#### Transcriptomic

Single-cell RNA-seq (scRNA-seq) for astrocytes was downloaded from a published study^69^. From 149,558 cells and 17,817 genes, we projected 4,292 metacells and 16,718 genes for AD pathologic diagnosis using *Metacell-2*^70^. Then, we performed differential expression testing for Alzheimer’s disease (AD) pathological diagnosis using *Seurat* and selected the top 280 differentially expressed genes (*p*-value adjusted < 0.1 and log_2_(fold change) > 0.2). Additionally, 25 genes that are either AD-risk or astrocyte-specific (*APOE*^12,45^, *CLU*^71^, *SORL1*^72^, *TREM2*^72^, *ABCA7*^72^, *PICALM*^72^, *BIN1*^48^, *PLD3*^72^, *CD33*^73^, *CD2AP*^72^, *EPHA1*^72^, *INPP5D*^72^, *FERMT2*^72^, *CR1*^72^, *PTK2B*^72^, *SLC24A4*^72^, *CASS4*^72^, *ACE*^73^, *SORCS1*^72^, *GAB2*^74^, *CDK5R1*^75^, *PRNP*^76^, *IL1RAP*^77^, *CNP*^53^, *TOMM40*^78^) were also selected.

#### Metabolomic

ROSMAP metabolomic data processed by *Metabolon HD4* was downloaded from *AD knowledge Portal* - backend (syn10235592). The dataset comprises a total of 1,055 biochemicals, 971 compounds of known identity (named biochemicals) and 84 compounds of unknown structural identity (unnamed biochemicals) for 514 brain samples. We selected 305 biochemicals that do not have any missingness in their samples.

#### Multi-view Dat

For early fusion, we merged transcriptomic and metabolomic data and achieved 2,286 metacells (samples) and 305 genes and 305 metabolites in each view. Metabolites were matched to the donors for metacells in transcriptomic data. For late fusion, we used full samples for each view whose AD pathologic diagnosis is known. 4,292 metacells and 2,286 samples for transcriptomics and metabolomics, respectively were used.

### Transcriptomics-Epigenomics (SEA-AD)

The *Seattle Alzheimer’s Disease Brain Cell Atlas* (SEA-AD) consortium studies deep molecular and cellular understanding of the early pathogenesis of AD^79^. By integrating neuropathological data, single-cell and spatial genomics, and longitudinal clinical metadata, SEA-AD serves as a unique resource for exploring the mechanisms underlying Alzheimer’s disease and related dementias.

#### Transcriptomic

scRNA-seq data for oligodendrocytes from middle temporal gyrus (111,194 cells) was downloaded from the *Registry of Open Data on AWS* as AnnData objects (h5ad format). 3,927 metacells and 17,368 genes were projected for cognitive status (dementia). We selected expressions for the 206 transcription factors that are known to co-bind or interact each other in oligodendrocytes^16^.

#### Epigenomic

Single-cell ATAC-seq (scATAC-seq) data for all nuclei from middle temporal gyrus was downloaded from the *Registry of Open Data on AWS* as AnnData objects (h5ad format). There were 35,925 cells and 218,882 peaks for oligodendrocytes. Among those peaks, we chose 702 peaks that are overlapped with the known oligodendrocyte-specific peaks^16^.

#### Multi-view Data

We merged the scRNA-seq and scATAC-seq datasets by cell. The 35,925 cells from the scATAC-seq data were then grouped according to the metacells they belonged to in the scRNA-seq data, and the average counts for each metacell were computed.

### Transcriptomics and Spatial Transcriptomics (SEA-AD)

#### Transcriptomic

scRNA-seq data for microglia was downloaded from *CellxGene*^80^ for a published study^79^. From 40,000 cells and 18,279 genes, we projected 1,009 metacells and 17,348 genes for AD progression score using *Metacell-2*. Then, we implemented differential expression testing for AD progression score using the Differential Gene Expression analysis based on the negative binomial distribution (*DESeq2*)^81^ and selected 400 differentially expressed genes (*p*-value adjusted < 0.05 and log_2_(fold change) > 0.75). Additionally, 28 genes that are either AD-risk or microglia-specific were identified. (*CSF1R*^82^, *TREM2*^72^, *SORL1*^72^, *CD68*^82^, *TNF*^83^, *CLEC7A*^82^, *IL6*^**108**^, *APOE*^12,45^, *C1QA*^82^, *C1QB*^82^, *C1QC*^82^, *SPP1*^82^, *MAPK8*^84^, *PTGS2*^85^, *VEGFA*^86^, *FOS*^82^, *CLU*^71^, *CR1*^72^, *PICALM*^72^, *CD33*^73^, *SORCS1*^72^, *GAB2*^74^, *CASS4*^72^, *CDK5R1*^82^, *PRNPL*^76^, *IL1RAP*^82^, *CNP*^53^, *TOMM40*^78^) were also selected.

#### Spatial Transcriptomics (MERFISH)

Astrocyte single-cell MERFISH data for the whole taxonomy collected from the middle temporal gyrus was obtained through the *Open Data Registry on AWS* as AnnData objects (h5ad format) for a published study^79^. From 1,321,191 cells and 135 genes, 8,693 metacells and 134 genes were projected for AD progression score using *Metacell-2*. 1,016 metacells were projected for astrocytes.

#### Multi-view Data

Among the 1,016 metacells in the MERFISH data, we randomly selected 1,009 metacells. The AD progression score was highly correlated between the scRNA-seq and MERFISH datasets (r = 0.986). For each dataset, we computed the mean AD progression score, which was highly correlated with the corresponding scores in both datasets (r = 0.997 for scRNA-seq and r = 0.996 for MERFISH).

### Adjusting for confounding

Since age and sex could act as confounders, we tested whether they were associated with both the exposure and the outcome. Specifically, we examined the following associations:

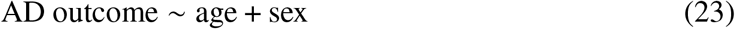

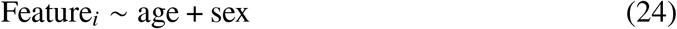

where *i* indexes each feature in two data views.

In the ROSMAP dataset used for binary classification (**Fig. 3**), sex was the only variable significantly associated with the outcome. Additionally, 230 out of 305 genes and 234 out of 305 metabolites were significantly associated with sex (FDR < 0.05). Therefore, we adjusted for sex by regressing out its confounding effect and used the adjusted data for prediction. All panels in **Fig. 3** have been updated to reflect the use of the adjusted input data.

In contrast, for the SEA-AD dataset used for continuous outcome prediction (**Fig. 5**), neither age nor sex showed a significant association with the outcome. Thus, they did not meet the criteria for confounding and were not adjusted for.

In the case of the multiome dataset used for binary classification (**Fig. 4**), since the samples were derived from metacells comprising cells from multiple donors, individual-level confounder adjustment was not applicable.

## Model Design

Let 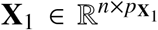 and 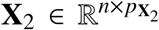 represent two data views, and **y**_1_, **y**_2_ ∈ ℝ^*n*^ be real-valued labels that represent the same disease phenotype corresponding to **X**_1_ and **X**_2_, respectively. COSIME inputs each data view into a deep neural network encoder (deep encoder), performs dimensionality reduction to learn *K*-dimensional features, and creates embeddings in an unsupervised fashion. Then, the Learnable Optimal Transport (LOT) aligns the distributions of two latent spaces in a joint latent space. We use the joint latent vector to classify or predict disease phenotypes.

### Deep Neural Network Encoders (Deep encoders)

COSIME is designed to process dual-view data using separate probabilistic deep encoders for each view. Each view *x*_1_ ∈ *X*_1_ and *x*_2_ ∈ *X*_2_ is passed through its own deep encoder to learn a latent representation. These encoders approximate the posterior distributions *p* (*z*_1_| *x*_1_) and *p* (*z*_2_ |*x*_2_), where *z*_1_ and *z*_2_ are the latent variables corresponding to views 1 and 2, respectively. COSIME utilizes backpropagation of Learnable Optimal Transport (LOT) to these deep neural networks, enabling the learning of latent features from multiple views, which is then used for predicting disease phenotypes.

The deep encoders output the mean (*μ*_*i*_) and log-standard deviation (log (*σ*_*i*_)) for the posterior distribution of each view. The encoder for each view consists of multiple fully connected layers with LeakyReLU activations. Batch Normalization is applied to stabilize learning and accelerate convergence, while Dropout is used as a regularization technique to mitigate overfitting. The final output of each deep encoder is a latent vector *z*_*i*_ that represents the compressed information from the corresponding input data of a view.

To regularize the learned posterior distributions and prevent overfitting, the model incorporates the Kullback-Leibler divergence (KLD) loss during the training of the deep encoders. This loss encourages the posterior distributions of each view to remain close to a standard Gaussian prior distribution *p* (*z*_*i*_) =𝒩 (0, *I*), which helps improve the ability of the model to generalize and prevents overfitting. The KLD loss for each view is computed as:

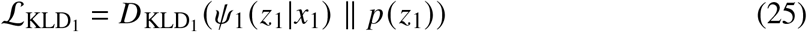

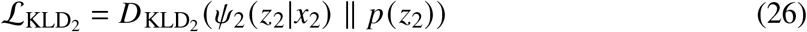

where *Ψ*_1_ and *Ψ*_2_ are the encoder functions for views *A* and *B*, respectively, 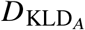and 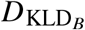 are the KLD measures for the latent representations of views *A* and *B*, respectively, and *p* (*z*_1_) and *p* (*z*_2_) are the prior distributions for the latent variables *z*_1_ and *z*_2_ corresponding to views *A* and *B*. The KLD loss for each view *i* is defined as:

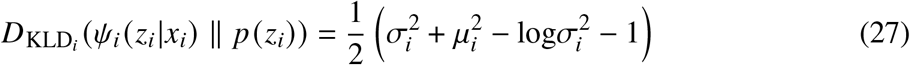

where *μ*_*i*_ and *σ*_*i*_ are the mean and standard deviation of the posterior distribution for view *i*.

### Learnable Optimal Transport (LOT)

COSIME utilizes Learnable Optimal Transport (LOT), a unique method designed to align the latent space distributions of two distinct data views. There are several key advantages of LOT.

First, unlike traditional optimal transport methods, Learnable Optimal Transport (LOT) treats the transport plan as a learnable parameter, optimized dynamically during training rather than relying on a fixed or pre-defined plan. This allows LOT to adapt more flexibly to complex data structures and align latent spaces in a joint latent space. By learning the transport plan iteratively, LOT can adjust its alignment strategy to the specific characteristics of the data, making it particularly effective for downstream tasks such as prediction and classification.

Second, LOT can scale effectively with large datasets. By incorporating mini-batch Sinkhorn^87^ updates, LOT accelerates the optimization process, reduces memory usage, and ensures efficient computation. This makes LOT particularly well-suited for high-dimensional data from multiple sources, enabling it to handle large-scale tasks without sacrificing performance.

Third, LOT integrates domain-specific regularization through the coupling matrix, which allows the model to incorporate prior knowledge about the relationship between the views. This is particularly beneficial when working with heterogeneous data in the multi-view analysis, where domain-specific relationships can enhance the quality of the learned latent space. We apply entropy regularization to stabilize the optimization process, ensuring smooth convergence and better-converging embeddings, which are crucial for downstream tasks.

#### Learnable Optimal Transport (LOT) Process

LOT treats the transport plan as a learnable parameter, which is iteratively updated through backpropagation. This enables LOT to adaptively align the latent spaces of different views in a shared representation space.

Both the source and target distributions in LOT are modeled as Gaussian distributions, each parameterized by their means (*μ*) and standard deviations (*σ*). The transport plan is learned by minimizing the pairwise transport cost between these distributions. The cost reflects the distance between their latent vectors, and the transport plan is updated during training to reduce this cost, aligning the distributions in a shared latent space.

The process involves the following steps:

#### 1. Step 1: Mini-Batching Processing

Before performing the computation, the source and target distributions are split into mini-batches to ensure efficient processing, especially for large datasets. Let the total number of source samples be *N*_src_ and the total number of target samples be *N*_tgt_. In the mini-batching procedure, we split the source distribution {*μ*_src_, *σ*_src_} and the target distribution {*μ*_tgt_, *σ*_tgt_} into smaller batches of size *B*_src_ and *B*_tgt_, respectively. Denote the *i*-th mini-batch of the source and target distributions as:

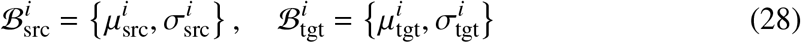

for *i* = 1, 2, …, *N*_src_ *B*_src_ and *j* = 1, 2, …, *N*_tgt_ *B*_tgt_, respectively. This mini-batch processing reduces memory consumption and allows the transport plan to be optimized iteratively.

Each mini-batch is processed separately, and the transport plan is updated incrementally based on these smaller subsets of the source and target distributions.

#### 2. Step 2: Pairwise Distance Computation

The pairwise distances between the means 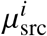 and 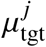 and the standard deviations 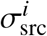 and 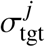 of the mini-batches are computed using the Euclidean distance:

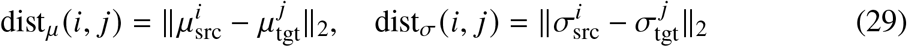

where *i* and *j* refer to the samples from the source and target mini-batches, respectively. These pairwise distances are used to construct the cost matrix for each mini-batch, which will then be used in the subsequent steps.

#### 3. Step 3: Cost Matrix Construction

After computing the pairwise distances, the total cost matrix for each mini-batch is constructed as follows:

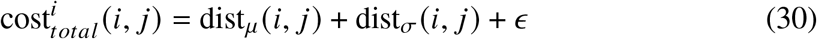

where *ϵ* is a small constant for numerical stability.

#### 4. Step 4: Sinkhorn Algorithm (Learnable Transport Plan)

The transport matrix **P**^*i*^ for each mini-batch is computed iteratively using the Sinkhorn algorithm. The transport plan is initialized as a learnable parameter, **P**^*i*^, where each element of **P**^*i*^ represents the transport probability between a sample from the source and a sample from the target. The transport plan is updated during each iteration to minimize the total transportation cost.

The update rule for the transport plan **P**^*i*^ (*i, j*) for each mini-batch is:

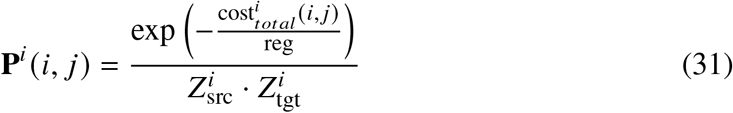

where 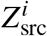 and 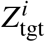 are the normalizing factors for the source and target distributions, respectively:

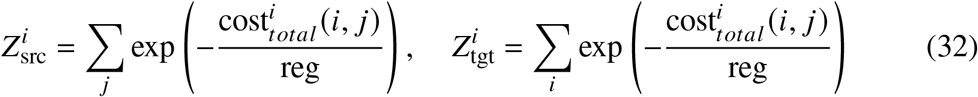

Here, the learnable transport plan allows gradients to flow through the transport matrix, enabling the optimization process using backpropagation.

#### Handling Different Sample Sizes

The Sinkhorn algorithm can effectively handle source and target distributions with differing sample sizes, making it particularly useful for COSIME late fusion. To ensure that transport plans remain valid, we apply normalization to the weights of the source and target distributions during each mini-batch update as follows:

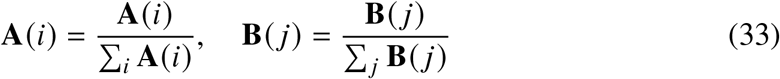

This adjustment ensures that the transport plan remains meaningful even when the source and target distributions contain different numbers of samples.

The Sinkhorn algorithm iteratively refines the transport plan **P**^*i*^ until convergence or until a predefined number of iterations.

#### 5. Step 5: Dual Variables and Marginal Constraints

Dual variables **u**^*i*^ and **v**^*i*^ are introduced to enforce the marginal constraints of the transport problem. These variables ensure that the transport plan satisfies the required marginal distributions for the source and target distributions. The dual variables are updated iteratively for each mini-batch:

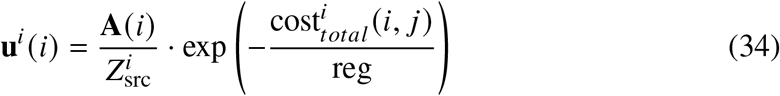

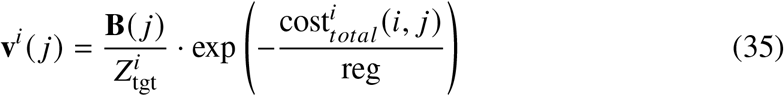

where **reg** denotes the entropic regularization parameter in Sinkhorn’s algorithm, and the dual variables enforce that the transport plan aligns with the marginal distributions of the source and target.

#### 6. Step 6: Optimizing the Transport Plan

During each mini-batch, the transport plan is optimized by performing backpropagation to minimize the total loss. This involves computing the gradients of the transport plan with respect to the loss function and updating the transport plan parameters via gradient descent.

#### 7. Step 7: Final LOT Loss Calculation

After optimizing the transport plan, the final LOT loss is computed as the sum of the product of the cost matrix and the transport plan:

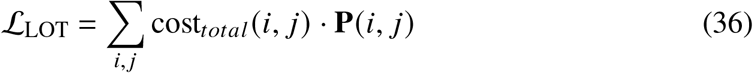

This loss function quantifies the minimum cost required to transform the source distribution into the target distribution, thus aligning the two views in a joint latent space.

#### Phenotype Prediction/Classification

COSIME employs the aligned latent representations from each view, obtained via separate probabilistic deep encoders. After the deep encoders generate the latent vectors for each view, LOT aligns these embeddings into a joint latent space. The model then uses these aligned embeddings for phenotype prediction.

Two fusion strategies are employed to combine the embeddings from the two views: early fusion and late fusion. Both fusion strategies are applied for binary outcome classification and continuous outcome prediction tasks.

#### Early Fusion

The fused representation is obtained by taking the mean of the representations *z*_1_ ∈ ℝ^*d*^ and *z*_2_ ∈ ℝ^*d*^ from each view:

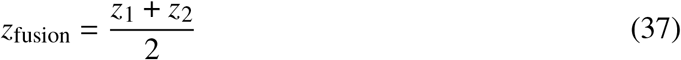

#### Late Fusion

The latent representations *z*_1_ ∈ ℝ^*d*^ and *z*_2_ ∈ ℝ^*d*^ from each view are vertically concatenated to form the fused representation *z*_fusion_:

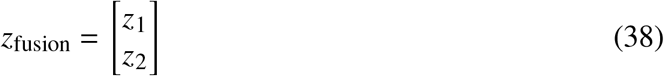

where *z*_fusion_ ∈ ℝ^2*d*^ is the fused vector obtained by stacking *z*_1_ and *z*_2_ vertically.

#### Binary Outcome Prediction

The fused representation *z*_fusion_ is then passed through a linear layer followed by a sigmoid activation function to compute the predicted probability 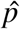:

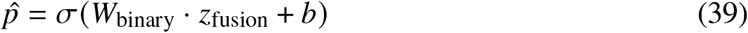

where *σ* (·) is the sigmoid function, and *W*_binary_ and *b* are the learned weights and bias parameters of the prediction layer.

#### Loss function

The **BCEWithLogitsLoss** is computed for the early fusion setup, which combines the sigmoid activation and binary cross-entropy in a single, more numerically stable operation. This loss function is defined as:

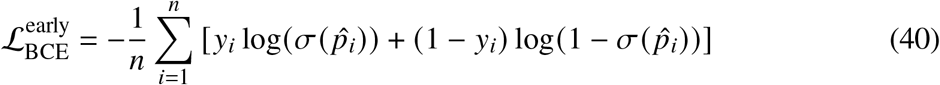

where 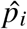 is the predicted probability of the *i*-th sample belonging to the positive class (i.e., the probability output by the model), and *y*_*i*_ is the true binary label for the *i*-th sample, where *y*_*i*_ ∈ {0, 1}.

#### Continuous Phenotype Prediction

For continuous phenotype prediction, the fused representation *z*_fusion_ is passed through a linear transformation:

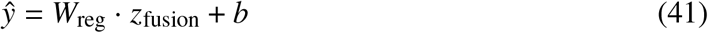

where *W*_reg_ and *b* are the learned weights and bias of the regression layer.

#### Loss for Regression

The Mean Squared Error (MSE) loss for continuous outcome prediction using a linear model is computed as:

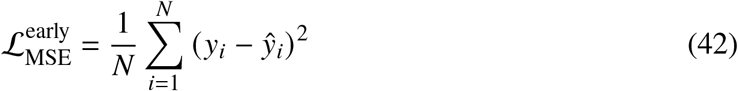

where ŷ_*i*_ is the predicted continuous value for the *i*-th sample, representing the model’s prediction, and *y*_*i*_ is the true continuous value for the *i*-th sample, which is the actual target value the model aims to predict. The loss measures the squared difference between the predicted and true values for each sample and averages this over all *N* samples.

#### Loss for Neural Network

A neural network (NN) model can be used for continuous outcome prediction, the output ŷ_*i*_ is predicted by the neural network, and the loss function remains MSE:

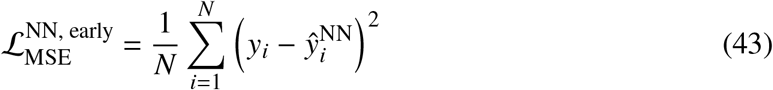

where 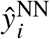 is the continuous value predicted by the neural network for the *i*-th sample, which is the output of the neural network model for that particular input, and *y*_*i*_ is the true continuous value for the *i*-th sample, which is the actual target value that the model is attempting to predict. The loss measures the squared difference between the predicted value and the true value for each sample, and the average squared error is computed over all *N* samples.

#### Prediction Loss

The prediction loss term ℒ_pred_ is defined based on the task at hand:

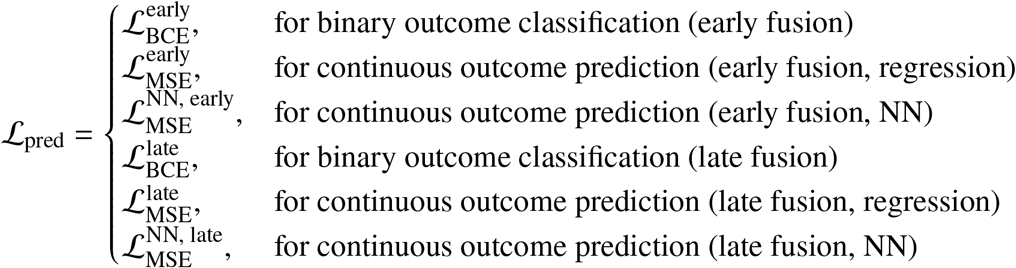

### Total Weighted Loss

During the training phase, the contributions of each loss term can vary in magnitude, depending on the relative scale of the individual losses. To balance the influence of each term on the total loss, we normalize the loss terms dynamically. This dynamic normalization ensures that no individual loss term has an outsized influence on the total loss, allowing for a more balanced optimization process.

We first compute the inverse of each loss term:

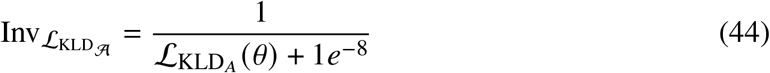

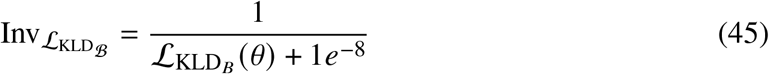

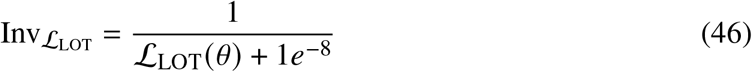

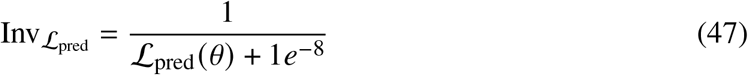

Next, we compute the total inverse magnitude:

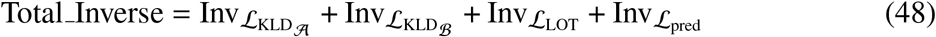

Now, we normalize the inverse magnitudes to get their relative adjusted weights:

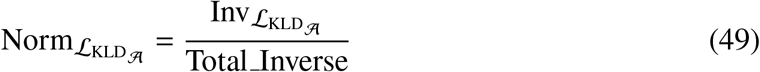

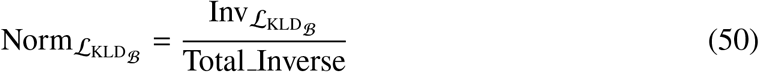

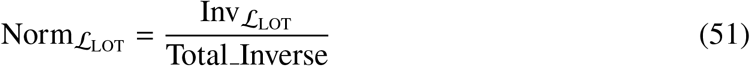

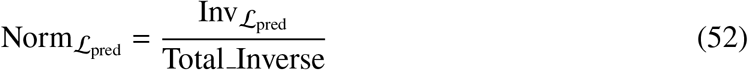

Finally, the total weighted loss ℒ_Total weighted_(*θ*) is computed as:

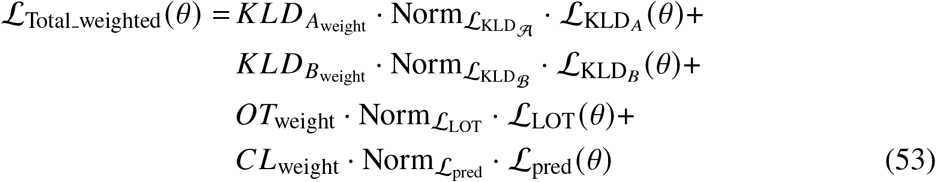

where 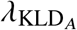, 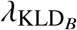, *λ*_LOT_, and *λ*_pred_ are the weights that control the importance of each normalized loss term, specifically 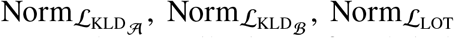, and 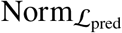, respectively. These normalized loss terms are the contributions of each individual loss term 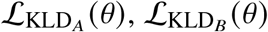, ℒ_LOT_ (*θ*), and ℒ _pred_ (*θ*)to the total loss, which are normalized to ensure that no single loss dominates the optimization process.

To optimize the model parameters, we minimize the total weighted loss:

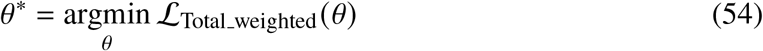

This method prevents overemphasis on any one loss component, allowing the model to effectively balance view alignments and prediction accuracy. It enables flexible adjustment of task importance, which is crucial when working with multi-view data, where different objectives (e.g., alignment, prediction) may compete for optimization.

## Model training process

The input data for each view was split into 75% for training and 25% for testing (holdout). 5-fold cross-validation was applied to the training set. The best-performing model was selected based on the minimum classification (or prediction) loss during the training phase and was then evaluated on the testing set. The **Algorithm 1: COSIME Model Training** is detailed in the **Supplementary Information**.

The prediction performance of the models was evaluated using holdout data with metrics such as Area Under the Receiver Operating Characteristic Curve (AUROC), Area Under the Precision-Recall Curve (AUPRC), and accuracy for binary outcome classification, and Mean Squared Error (MSE) for continuous outcome prediction. The performance metrics were compared across different methods using the mean and ±1.96 times the standard deviation. For the best-trained models across the different multi-view datasets, we computed feature importance for each data view and interaction values both within and across views.

### Hyperparameter tuning

We tuned several hyperparameters such as the learning rate, learning rate decay (gamma), loss weights, dropout rate, joint latent space dimensionality, early stopping patience, minimum change for early stopping, and weight decay in the optimizer. Hyperparameter optimization was performed using *Ray Tune*^88^ with Distributed Asynchronous Hyperparameter Optimization (*Hyperopt*)^89^. The Adam optimizer was employed to minimize the total loss, with a maximum of 300 epochs. Early stopping was used to prevent overfitting, based on performance improvements on the validation set in the training phase.

### Optimization and Loss Minimization

During the training phase, the model updates the hyper-parameters by minimizing the total loss. The optimization process involves computing gradients with respect to the total loss function and updating the parameters using gradient descent or its variants.

By minimizing the total loss, COSIME simultaneously learns to:

- Regularize the posterior distributions of the latent representations for each view through the KLD losses,
- Align the embeddings from both views into a shared latent space using LOT,
- Predict or classify the disease phenotype by minimizing classification (or prediction) loss.

### Evaluating the Performance of the Model Predictions

To evaluate the performance of the best model selected in the training phase for both binary outcome classification and continuous outcome prediction, several standard metrics are employed:

### Binary Outcome Classification

#### 1. Area Under the Receiver Operating Characteristic Curve (AUROC)

AUROC measures the ability of the model to discriminate between the positive and negative classes. It is calculated by assessing the True Positive Rate (TPR) against the False Positive Rate (FPR) at various threshold settings. The AUROC value ranges from 0 to 1, where 1 indicates perfect classification, and 0.5 is the expected performance of a random classifier. The formula for AUROC is:

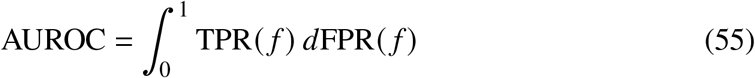

where TPR (*f*) and FPR (*f*) are the True Positive Rate and False Positive Rate at each decision threshold *f*.

#### 2. Area Under the Precision-Recall Curve (AUPRC)

AUPRC is another useful metric for evaluating binary classification performance, particularly when dealing with imbalanced datasets. It measures the area under the curve created by plotting precision against recall at various threshold settings. The formula for AUPRC is:

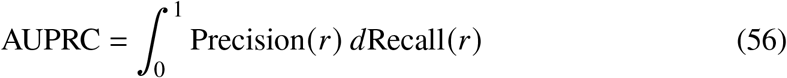

where Precision(*r*) and Recall(*r*) are the precision and recall at each decision threshold *r*.

#### 3. Accuracy

Accuracy is the proportion of correctly classified instances (both true positives and true negatives) out of the total number of instances. It is calculated as:

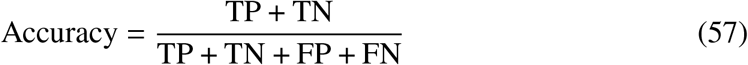

where TP is True Positives, TN is True Negatives, FP is False Positives, and FN is False Negatives.

### Continuous Outcome Prediction

#### Mean Squared Error (MSE)

MSE measures the average squared difference between the true values and the predicted values. It is a common loss function for regression tasks and is used to quantify the error in continuous prediction tasks. The formula for MSE is:

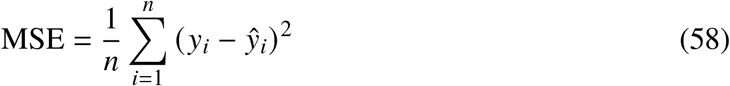

where *y*_*i*_ is the true continuous label for the *i*-th sample, ŷ_*i*_ is the predicted value for the *i*-th sample, and *n* is the total number of samples.

These evaluation metrics assess the effectiveness of the COSIME model in both binary and continuous phenotype prediction tasks. Higher AUROC and AUPRC values, along with higher accuracy and lower MSE, indicate better model performance.

## FI and feature interaction

COSIME introduces an advanced approach for quantifying both FI and feature interactions in predictive modeling, building upon the Shapley-Taylor framework^8^. By using Monte Carlo sampling and a batching process, COSIME efficiently approximates Shapley values and Shapley-Taylor indices, making it highly scalable and interpretable for large, high-dimensional datasets while maintaining memory efficiency. This approach ensures COSIME is well-suited for complex real-world applications.

FI is simply the difference between a model’s prediction when using all features and its prediction when one specific feature is left out. Feature interaction refers to the additional effect on the prediction when two features are used together, beyond the sum of their individual effects. Note that pairwise feature interaction is different from traditional correlation between two features, which describes their relationship in the input data but does not reflect their influence on the model’s prediction.

COSIME employs Monte Carlo sampling to efficiently approximate both Shapley values and feature interactions. This approach notably enhances its scalability, interpretability, and flexibility, especially when handling large datasets and high-dimensional feature spaces. Instead of relying on exact calculations, which become computationally expensive as the number of features or data points increases, Monte Carlo sampling allows COSIME to estimate Shapley values and Shapley-Taylor indices efficiently. This method also offers precise and interpretable calculations of pairwise feature interactions by directly simulating feature combinations. By estimating these interactions in this way, COSIME avoids complex approximations or indirect assumptions, providing clearer insights into how combinations of features jointly affect predictions. Furthermore, COSIME is better equipped to handle complex feature interactions and nonlinear relationships more effectively.

Additionally, COSIME is memory-efficient due to its batching process, which breaks the data into smaller subsets to minimize memory usage. This enables COSIME to handle large datasets on systems with limited memory, making it suitable for real-world applications with resource constraints. The batching process ensures that COSIME can scale efficiently while maintaining performance across high-dimensional feature spaces.

COSIME can also calculate signed feature interactions, which indicate whether pairs of features interact synergistically (positive interaction) or antagonistically (negative interaction). This is achieved by directly simulating feature combinations during the Monte Carlo sampling process, allowing COSIME to estimate both the magnitude and direction of interactions. The sign of an interaction is determined by comparing the joint effect of the features to their individual effects, revealing whether their combined influence on the prediction. This enhancement adds a important layer of interpretability, enabling users to understand not only the strength but also the directionality of feature interactions.

Finally, COSIME allows users to customize the Monte Carlo sample size and maximum memory usage, providing flexibility in the computational process. When the memory usage is specified, COSIME automatically determines the optimal batch size, making the method both user-friendly and adaptable to different system configurations. The details of **Algorithm 2: COSIME Feature Importance and Interaction Computation** are provided in the **Supplementary Information**.

### Shapley Value (Feature Importance Computation)

The Shapley value *ϕ*_*i*_ for a given feature *i* is computed by considering the marginal contribution of feature *i* across all possible combinations of other features. This is approximated using Monte Carlo sampling over multiple iterations. The Shapley value for each feature *i* and each sample *s* is computed as:

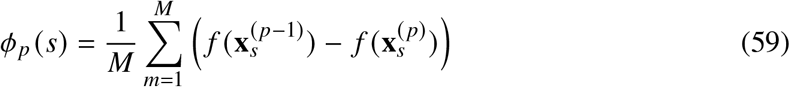

where:

- *f* (**x**_*s*_) represents the prediction using all features,
- 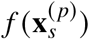represents the prediction when feature *p* is masked,
- 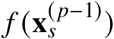represents the prediction when all features except for feature *p* are masked.

The Shapley values for all features are stored in a matrix 𝒮∈ ℝ^*N*×*F*^, where *N* is the number of samples and *F* is the number of features.

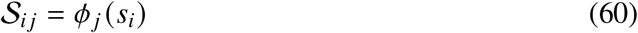

where *i* indexes the *i*-th sample (*i* ∈ {1, 2, …, *N* }) and *j* indexes the *j*-th feature (*j* ∈ {1, 2, …, *F* }), and *ϕ* _*j*_ (*s*_*i*_) represents the Shapley value for feature *j* with respect to sample *i*.

### Feature Interaction Effects

The interaction effect between two features *p* and *q* quantifies the joint contribution of features *p* and *q* compared to when they are individually absent. The interaction effect ℐ_*pq*_ is computed as:

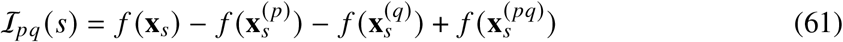

where:

- *f* (**x**_*s*_): the prediction of the model using all features,
- 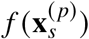and 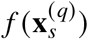: the predictions of the model with features *p* and *q* individually masked,
- 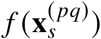: the prediction of the model when both features *p* and *q* are masked.

For pairwise interactions, the interaction effect is averaged over *M* Monte Carlo iterations:

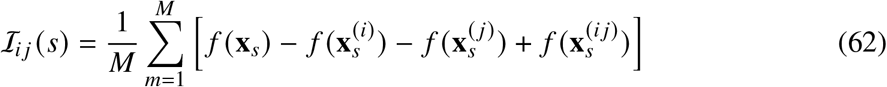

The interaction effects for all pairs of features are stored in the interaction matrix ℐ ∈ ℝ^*F*×*F*^, where each entry ℐ_*i j*_ represents the interaction between features *i* and *j*.

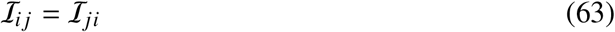

### Monte Carlo Approach and Interaction Algorithm

Given the model *f*, input data *X* ∈ ℝ^*N*×*F*^ (where *N* is the number of samples and *F* is the number of features), and target labels *y* ∈ ℝ^*N*^, we compute the Shapley values and interaction effects as follows:

- **Shapley Values**: For each feature *i*, compute the marginal contribution to each sample over *M* Monte Carlo iterations. Store the results in the matrix 𝒮.
- **Interaction Effects**: For each pair of features (*i, j*), compute the interaction effect using the formula above, averaging over *M* Monte Carlo iterations. Store the results in the interaction matrix ℐ.

The algorithm processes the data in batches to manage memory efficiently. If the batch size is not provided, it is dynamically computed based on the available memory. The number of batches *B* is given by:

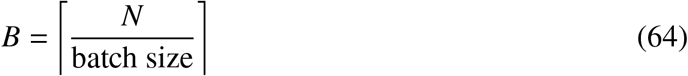

The batch processing ensures that we do not exceed the maximum memory usage during computation. Considering the available resources and estimated computing time, we used 100 Monte Carlo iterations for the simulated multi-view data and 10 iterations for the real multi-view data. For each multi-view dataset, the test (holdout) data was used.

### Memory Management

To ensure that the memory usage remains within a specified limit, we compute the memory required for each batch as:

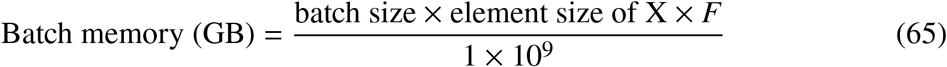

If the calculated batch memory exceeds the available memory, the batch size is adjusted accordingly. The dynamic batch size calculation ensures that the computation does not exceed the memory limit.

### Output

The Shapley values are stored in a matrix 𝒮 with dimensions *N* × *F*, and the interaction effects are stored in a matrix ℐ with dimensions *F* × *F*. These matrices represent the individual contributions of each feature and their pairwise interactions in the prediction process.

In summary, our innovative approach provides a comprehensive methodology for assessing both individual feature importance (Shapley values) and pairwise feature interactions in machine learning models. By leveraging these tools, we enable a deeper understanding of how features individually and collectively influence model predictions. The computational framework we present, along with the accompanying code, facilitates the application of this methodology to various prediction tasks, enhancing model interpretability and providing valuable insights into feature relationships.

## Downstream analyses

### Gene Enrichment Analysis

The function `enrichGO` from the `clusterProfiler`^90^ R package, a universal enrichment tool for interpreting omics data, was used to perform enrichment analyses for the 20 prioritized genes by absolute feature importance (FI) values in each view of real multi-view datasets. The top 10 categories were displayed by log_10_(FDR) (**Fig. 3b** and **Supplementary Fig. 1**).

### Metabolite Enrichment Analysis

`MetaboAnalyst` 6.0^91^ was used to implement a metabolite set enrichment analysis for 20 prioritized metabolites by absolute feature FI values in the metabolomic data for classifying cognitive diagnosis for Alzheimer’s disease (**Fig. 3c**).

### Interaction Network

For the top 50 across-view interaction values, the edges and nodes for the interacting genes (and genes and metabolites) were formatted in R, and the network was generated using `Cytoscape 3.10.3`^92^ (**Fig. 3e** and **Fig. 5d**).

### Spatial mapping and statistical tests

Spatial expression plots for *RYR3* and *DGKG* were generated in Python, employing matplotlib and seaborn (**Fig. 5c**). To assess whether there are significant spatial differences between cells with high FI values and low feature importance values for each gene, the following steps were taken:

- Compute the midpoint of the bounding box: It is calculated by finding the midpoint of the bounding box defined by the *x* and *y* coordinates. Specifically, the center is determined by averaging the maximum and minimum values of both the *x* and *y* coordinates:

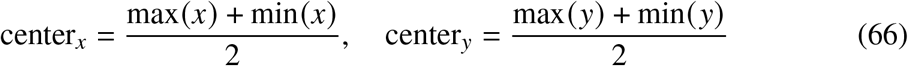

This central point serves as the reference from which the distances of each point are measured.

- Compute the Euclidean distance from each point to the midpoint of the bounding box: Once the midpoint of the bounding box is defined, the Euclidean distance from each point (*x*_*i*_, *y*_*i*_) to the computed midpoint of the bounding box (center_*x*_, center_*y*_) is calculated using the formula:

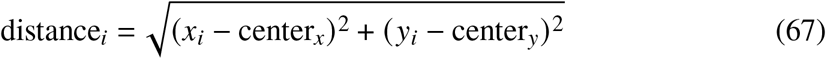

This step computes a scalar value for each point that quantifies how far each point is from the center.

- Perform a two-sample two-tailed t-test: The dataset is divided into two groups based on the feature importance:

- Group 1: Points with FI values greater than or equal to the 75th percentile (i.e., the top 25%).
- Group 2: Points with FI values below the 75th percentile.

We performed a two-sample two-tailed t-test to evaluate whether the distances from the midpoint of the bounding box differed between the top 25% of cells with the highest feature importance values and the remaining 75% of cells. The null hypothesis (*H*_0_) for the two-tailed t-test is:

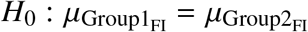

where 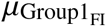 and 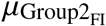 represent the means of the distances for the high and low FI groups, respectively. The alternative hypothesis is:

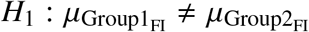

The test statistic *t* and the associated *p*–value are calculated to determine whether the difference in means is statistically significant.

Similarly, the spatial distribution of across-view interaction values was visualized using the same tools (**Fig. 5e**). The following steps were taken to generate the spatial distribution of across-view interaction values and assess whether there are significant spatial differences between cells with positive feature interaction values and negative feature interaction values, the subsequent steps were carried out:

- Distance calculation: The distance of each cell from the center of the tissue sample was calculated using its spatial coordinates (*X, Y*). This distance quantifies how far each cell is from the central point of the tissue section. The Euclidean distance from the center is computed as:

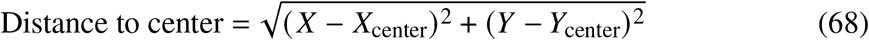

where *X*_center_ and *Y*_center_ are the coordinates of the center of the tissue.
- Cell-level interaction score: For each cell, the interaction score was computed using the MERFISH gene expression levels and pairwise across-view interaction values. Let 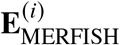 denote the gene expression vector for cell *i*, which is of length *G*_MERFISH_. The interaction matrix **M** has dimensions *G*_MERFISH_ × *G*_scRNA-seq_, where each element 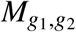 represents the pairwise interaction between MERFISH gene *g*_1_ and scRNA-seq gene *g*_2_. The cell-level interaction score *S*^(*i*)^ for each cell *i* was calculated as the dot product of the gene expression vector for that cell and the interaction matrix:

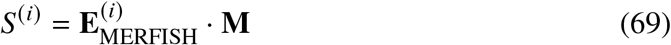

In expanded form, the score for cell *i* with respect to scRNA-seq gene *j* is:

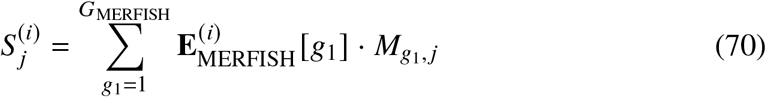

This score quantifies the interaction between the MERFISH gene expression profile of cell *i* and the scRNA-seq gene expression profile, enabling grouping of cells based on their interaction scores. Cells were then grouped based on the sign of their interaction scores: those with positive scores were assigned to the Positive group, while those with negative scores were assigned to the Negative group. This grouping allowed for further comparison of spatial distributions and other characteristics between the two groups.

- Perform a two-sample two-tailed t-test: To determine whether there were significant differences in the distances of cells from the center between the two groups, an independent two-sample *t*-test was performed. The hypotheses tested were:

- Null hypothesis (*H*_0_): The mean distance from the center of the Positive and Negative groups are equal, i.e.,

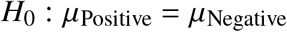
- Alternative hypothesis (*H*_1_): The mean distance from the center of the Positive and Negative groups are not equal, i.e.,

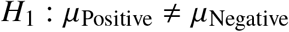

- Perform a two-tailed F-test: In addition to comparing the means, an *F*-test was performed to compare the variances of the two groups. The hypotheses tested were:

- Null hypothesis (*H*_0_): The variances of the Positive and Negative groups are equal, i.e.,

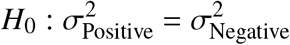
- Alternative hypothesis (*H*_1_): The variances of the Positive and Negative groups are not equal, i.e.,

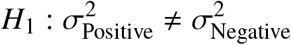

## Benchmarking methods

### Cooperative Learning

Cooperative learning is a framework for fitting machine learning models to multi-view data^3^. It presents a loss function that minimizes the prediction error across all views, while encouraging the predictions from the views to agree. This framework with linear regression, which is what we use for benchmarking.

We implemented the Cooperative Learning framework using the `multiview` R package. Following the examples provided on the Cooperative Learning GitHub repository, we preprocessed the input data using the *preProcess* function to center and scale it. For model evaluation, we conducted 5-fold cross-validation using the `cv.multiview` function, with a maximum of 300 iterations. For binary outcome classification, we set `family = binomial()` and type.measure = “class”. For continuous outcome prediction, we used the default `family = gaussian()` and `type.measure = “mse”`. The hyperparameter values for the regularization parameter were chosen from the set {0, 0.2, 0.4, 0.6, 0.8, 1}, while the lambda values were left at their default settings.

### Data Integration Analysis for Biomarker discovery using Latent components (DIABLO)

Data Integration Analysis for Biomarker discovery using Latent components (DIABLO) is a supervised extension of canonical correlation analysis^4^. It aims to identify linear combinations of variables from each dataset that maximize the correlation between datasets.

We implemented DIABLO using the `mixOmics` R package. Following the tutorial on the mixOmics website, we created a data block using the matrix function. For model training, we performed 5-fold cross-validation using the `block.splsda` function. The number of variables was tested within the set {10, 20, 30, 40, 50}.

### Multi-Omics Factor Analysis v2 (MOFA+)

Multi-Omics Factor Analysis v2 (MOFA+) is an unsupervised matrix decomposition method that can be viewed as a generalization of sparse principal components analysis.

We used the `MOFA2` R package to obtain latent representations. Following the tutorial on the MOFA+ downstream analysis in R, we trained a MOFA model using the `run mofa` function. To project unseen data from a new view into the latent space *Z*_*i*_, we computed the pseudo-inverse of the weight matrix *W*_*i*_. The latent representations *Z*_*i*_ for all views were then concatenated. For prediction, we applied logistic regression for binary outcomes and linear regression for continuous outcomes. To classify and predict new, unseen samples, we used the concatenated latent representation matrix *Z* in PyTorch (Python 3.10). We evaluated different latent factor dimensions from the set {1, 5, 10, 15}.

cha

### Aligned Optimal Transport

Aligned optimal transport (Aligned OT)^93^ serves as a baseline OT loss, where the OT matrix is assumed to be diagonal. Therefore, the final cost is the sum of the diagonal elements of the cost matrix, which we use in place of the LOT loss for our benchmark. Aligned OT thus captures only the alignment between the two latent spaces.

To compute this loss, we used the Aligned OT implementation from the Python Optimal Transport (POT)^93^ library. For variability analysis, we quantified the dispersion of cost values by computing the standard deviation of the cost matrices in both source and target modalities relative to their means.

### Single-cell Multi-omic Integration with Optimal Transport

The Single-cell Multi-omic Integration with Optimal Transport (SCOT)^94^ OT loss is more complex leverages Gromov-Wasserstein transport to capture structural relationships between the source and target cost matrices. The resulting transport plan is combined with the cost matrices through element-wise multiplication and summation to yield the final OT loss. This approach uses Euclidean distances to define costs and seeks to minimize distances between matched representations across modalities.

We implemented the OT loss computation by referring to SCOT methods. To assess variability in transport costs, we again measured the standard deviation of the cost matrices with respect to their means in each modality.

## Data Availability

Simulated data and all data supporting the results are included in the Supplementary Data 1–9 and are publicly available on GitHub (https://github.com/daifengwanglab/COSIME). ROSMAP scRNA-seq was sourced from a website (compbio.mit.edu/ad aging brain). Metabolomic data can be downloaded from the AD Knowledge Portal - backend (syn10235592). SEA-AD scATAC-seq, scRNA-seq, and MERFISH data were obtained from SEA-AD: Seattle Alzheimer’s Disease Brain Cell Atlas (cellxgene.cziscience.com/collections/…).

## Code Availability

The entire COSIME code is implemented in Python and available via Zenodo at https://doi.org/10.5281/zenodo.15428769^95^. The COSIME Python package and examples can be accessed publicly at https://github.com/daifengwanglab/COSIME.

## Acknowledgements

J.J.C. was supported by National Institutes of Health (NIH) grants under funding codes R21NS128761, RF1MH128695, R01AG067025, and P50HD105353, all awarded to D.W.; the National Science Foundation CAREER Award 2144475, also to D.W.; the Wisconsin Distinguished Graduate Fellowship (WDGF) from the Waisman Center; and an NIH grant under funding code RF1AG054047 awarded to C.D.E. N.C.K. and T.G. were supported by the same NIH and NSF grants awarded to D.W. (R21NS128761, RF1MH128695, R01AG067025, P50HD105353, and NSF CAREER Award 2144475). T.L. was supported by NIH grant RF1AG054047 awarded to C.D.E.

## Author Information

### Authors and Affiliations

**Department of Population Health Sciences, University of Wisconsin-Madison, Madison, WI 53726, USA**

Jerome J. Choi, Corinne D. Engelman, and Tianyuan Lu

**Waisman Center, University of Wisconsin-Madison, Madison, WI 53705, USA**

Jerome J. Choi, Noah Cohen Kalafut, and Daifeng Wang

**Department of Computer Sciences, University of Wisconsin-Madison, Madison, WI 5706, USA**

Noah Cohen Kalafut and Daifeng Wang

**Department of Biostatistics and Medical Informatics, University of Wisconsin-Madison, Madison, WI 53076, USA**

Tim Gruenloh and Daifeng Wang

### Contributions

Conceptualization, J.J.C. and D.W.; Methodology, J.J.C. and N.C.K.; Formal Analysis, J.J.C. and T.G.; Investigation, J.J.C., T.L., and D.W.; Writing–Original Draft, J.J.C.; Writing–Review and Edit, J.J.C., N.C.K., T.G., C.D.E., T.L., and D.W.; Supervision, T.L. and D.W.; Funding Acquisition, C.D.E. and D.W..

### Corresponding Authors

Correspondence to Daifeng Wang.

