## Supplementary Materials for "Cooperative multi-view integration with Scalable and Interpretable Model Explainer"

### Supplementary information

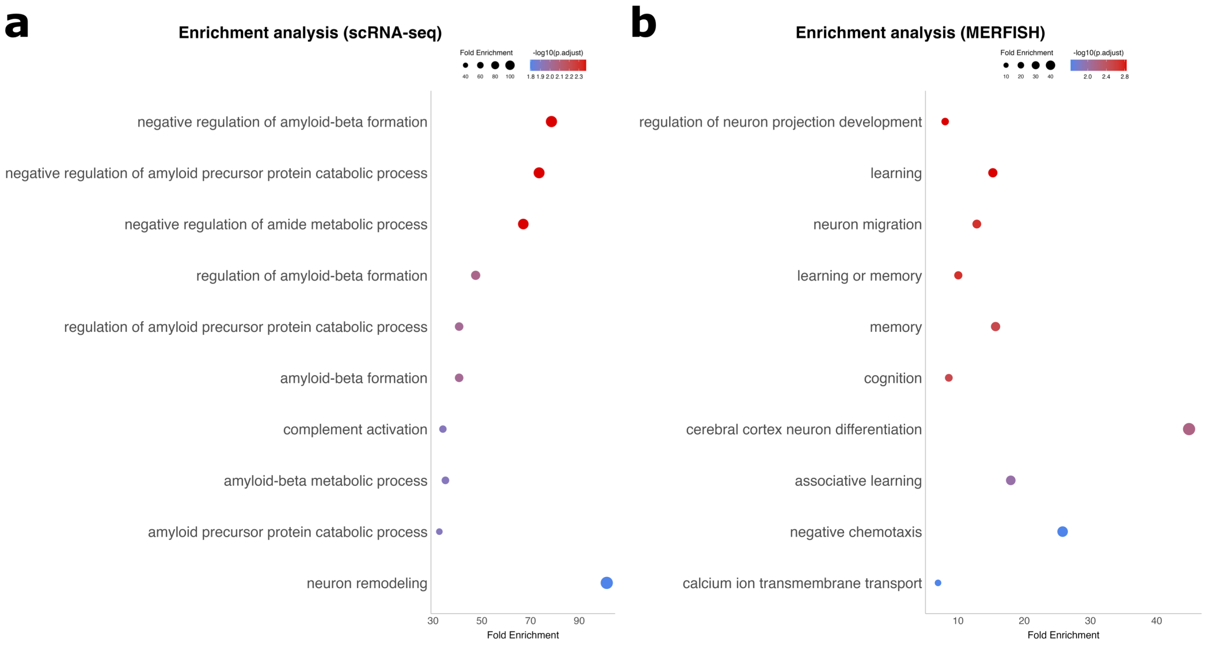


**Supplementary Fig. 1: Enrichment analyses prioritized for each view.**
**a** The top 20 prioritized genes (absolute feature importance values) in microglia **b** The top 20 prioritized genes (absolute feature importance values)in astrocytes.

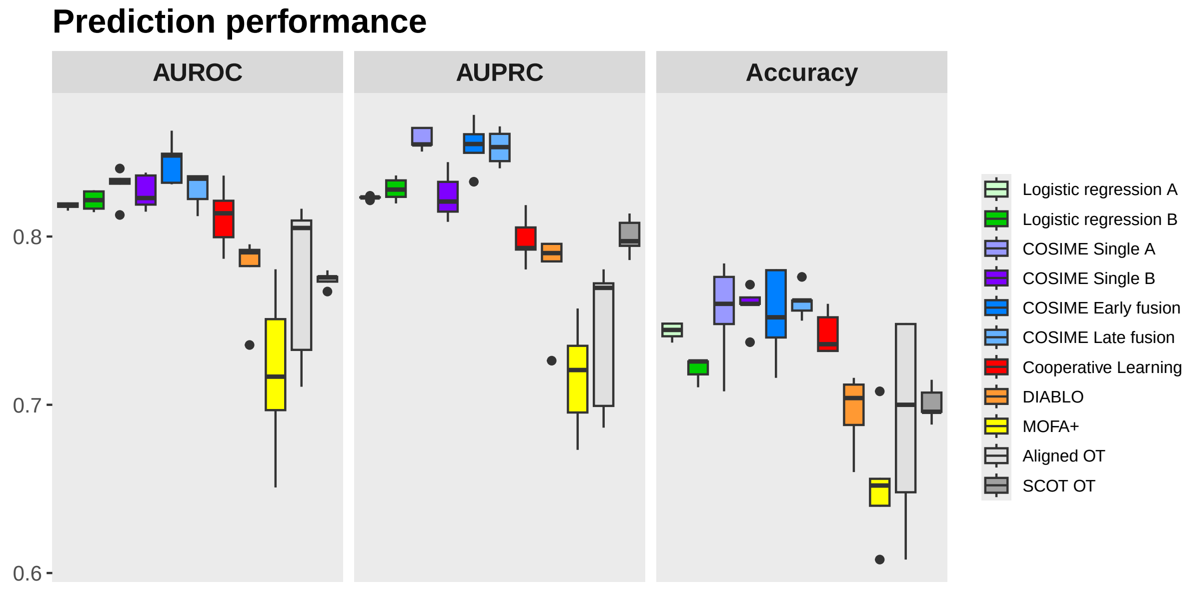


**Supplementary Fig. 2: Binary classification (Simulated data - high signal).**

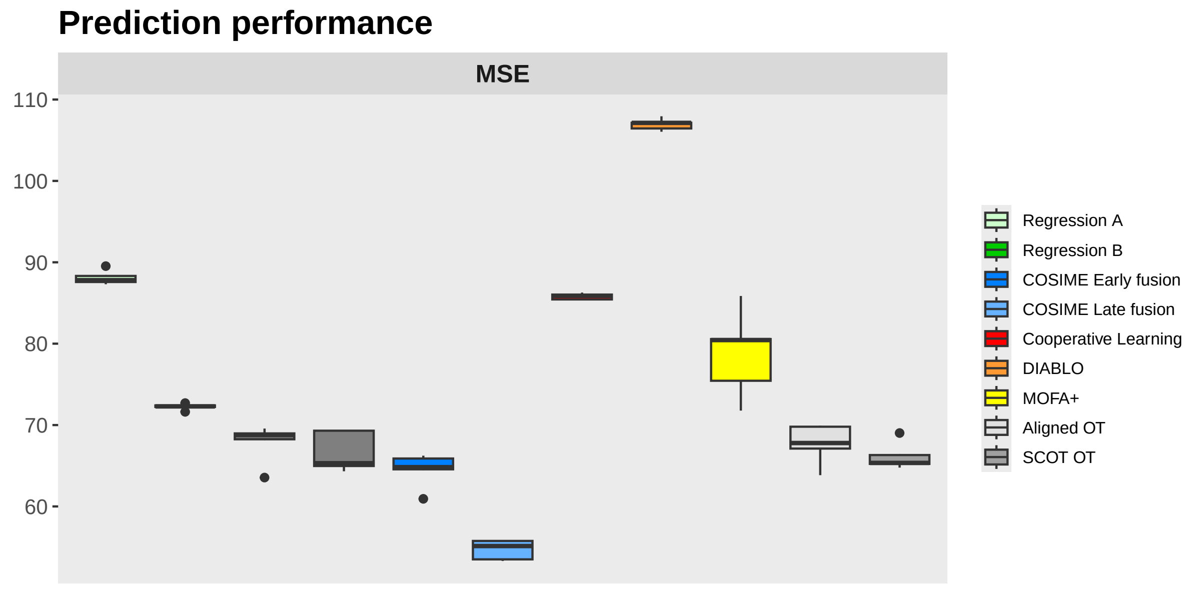


**Supplementary Fig. 3: Continuous outcome prediction (Simulated data - high signal).**

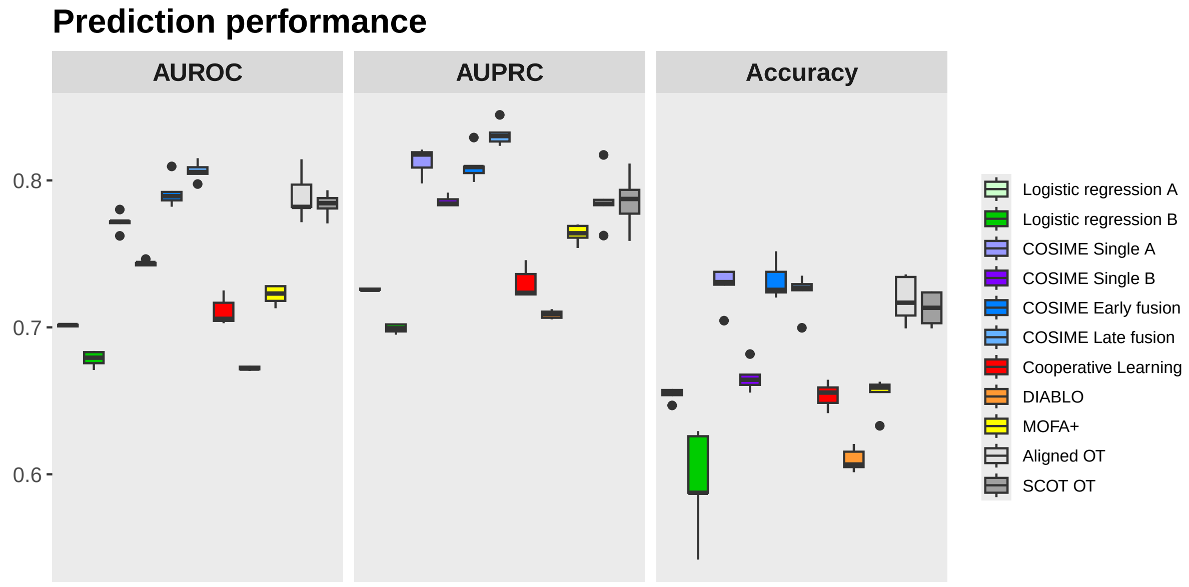


**Supplementary Fig. 4: Classifying cognitive diagnosis for Alzheimer's disease from transcriptomic (astrocytes) and metabolomic data.**

#### Supplementary Note 1: Performance across different data distributions

COSIME does not assume direct correspondence or equivalence between fundamentally different data types. Instead, each modality is first encoded into its own modality-specific latent representation through a deep encoder that preserves modality-specific structure. LOT then aligns these latent distributions rather than the raw features. Importantly, LOT does not enforce rigid alignment; it learns a soft and flexible transport plan that minimizes the cost of moving probability mass from one latent distribution to another. This transport plan is dynamically optimized during training and can reflect view-specific regularization, allowing the model to preserve differences in modality-specific representations while still learning shared patterns relevant to the outcome. In other words, LOT preserves the inherent geometry of each view's latent space and aligns them only to the extent needed for phenotype prediction. This design allows COSIME to account for the within-view heterogeneity while discovering shared patterns across views.

The empirical results (**Figs. 3–5**) demonstrate that COSIME maintains high predictive performance across diverse modality combinations—including transcriptomics, metabolomics, ATAC-seq, and spatial transcriptomics—supporting the robustness and biological validity of the learned shared space.

From a biological perspective, different omics data types can capture complementary views of the same underlying biological system, such as gene regulatory programs, cell-type identity, or disease status. These shared biological processes motivate the integration of distinct data modalities into a common latent space that emphasizes biologically meaningful variation.

The different distributions in the real data are presented in **Supplementary Figs. 5 to 7**, with four randomly selected features shown for each data view.

**Supplementary Fig. 5** presents distributions of four randomly selected features from the ROSMAP data for two different data views. **Fig. 5a** displays gene expression distributions from the ROSMAP transcriptomic data, where individual cells were aggregated into meta cells. This aggregation reduces sparsity and variability, resulting in smoother, continuous distributions that retain the characteristic right skew and overdispersion of the original single-cell data. **Fig. 5b** shows distributions of selected metabolomic features from the same cohort. These are inherently continuous and also right-skewed, with density shapes and ranges varying by metabolite. The observed variation reflects both biological differences in metabolite abundance.

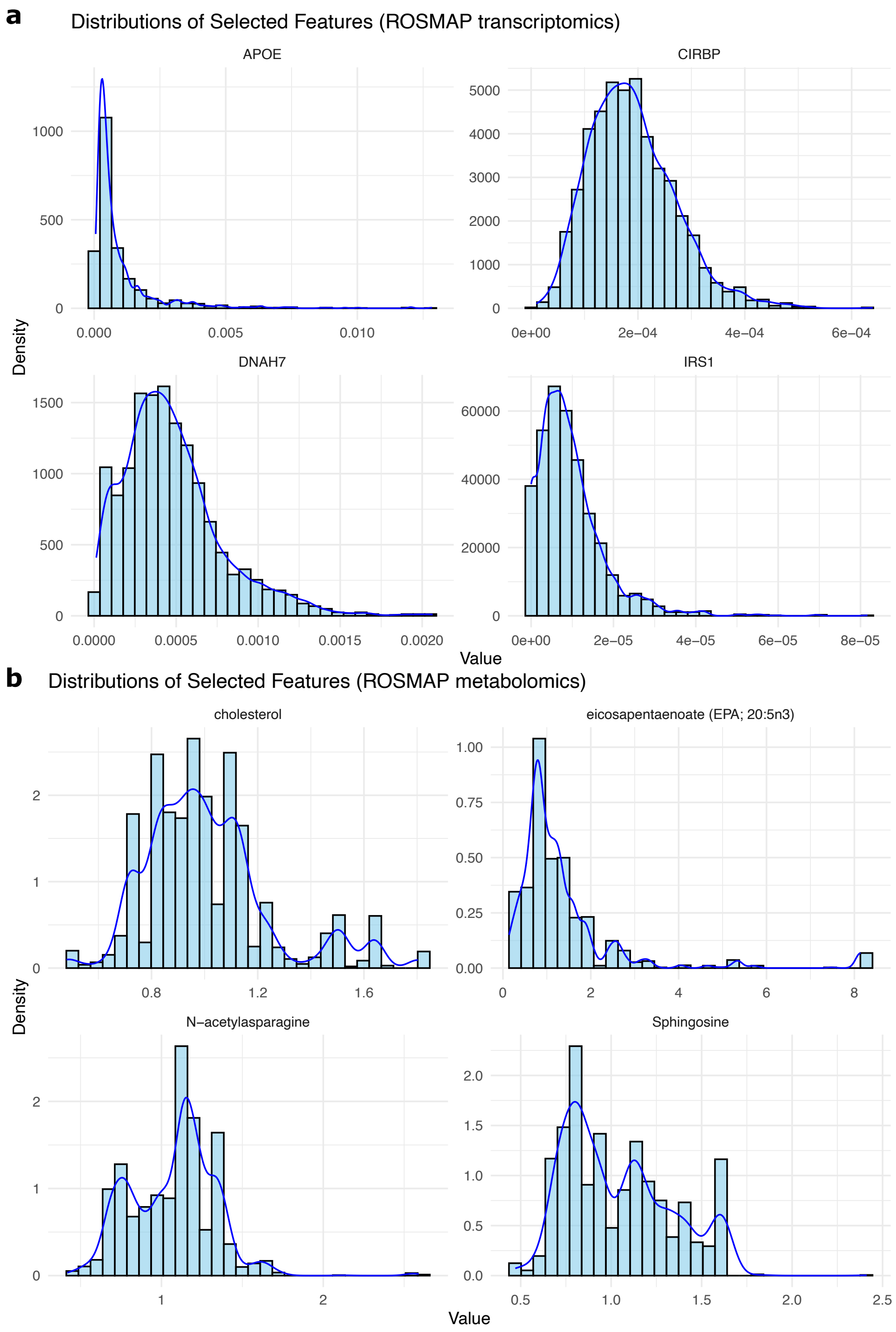


**Supplementary Fig. 5: Distributions of transcriptomic and metabolomic data.**
**a** Distributions of four gene expressions (Astrocytes) **b** Distributions of four metabolite levels (metabolomics)

**Supplementary Fig. 6** illustrates distributions of four randomly selected features from the SEA-AD data for two different data views. **Supplementary Fig. 6a** presents distributions of selected chromatin accessibility peaks from scATAC-seq data. These distributions are continuous, sparse, and right-skewed, with most values concentrated near zero and some features showing broader variability. The sparsity and shape reflect the underlying binary nature of accessibility as well as the effects of normalization and aggregation across meta cells. **Supplementary Fig. 6b** visualizes distributions of gene expression features from scRNA-seq data, also derived from meta cells. While raw single-cell RNA-seq data typically follow a NB distribution due to their discrete, overdispersed nature, aggregation into meta cells reduces sparsity and smooths the distributions. As a result, the displayed features retain NB-like characteristics such as right skew and overdispersion but no longer strictly follow a discrete NB form.

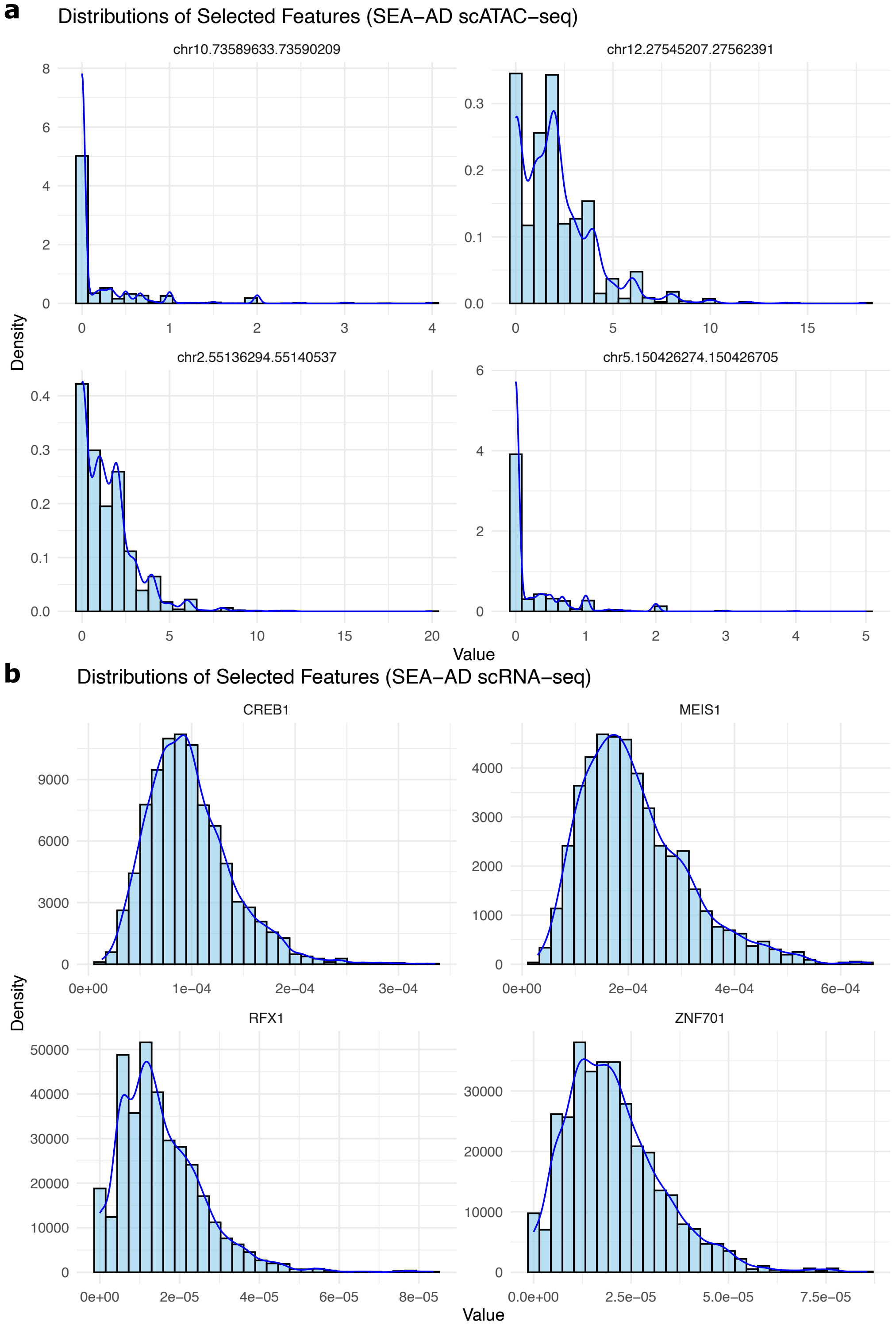


**Supplementary Fig. 6: Distributions of oligodendrocytes scRNA-seq and scATAC-seq data.**
**a** Distributions of four gene expressions **b** Distributions of four peaks (counts).

**Supplementary Fig. 7** depicts distributions of four randomly selected features from the SEA-AD dataset for two different cell types and data views. **Supplementary Fig. 7a** illustrates gene expression distributions from scRNA-seq microglia data aggregated into meta cells. While raw single-cell RNA-seq data typically follow a NB distribution due to their discrete, overdispersed, and zero-inflated nature, the use of meta cells reduces sparsity and variability. As a result, the displayed distributions are smoother and continuous, retaining NB-like characteristics such as right skew and overdispersion without strictly following a discrete NB form. **Supplementary Fig. 7b** demostrates distributions from the MERFISH spatial transcriptomic data in astrocytes. These distributions are continuous and right-skewed, with variability across genes reflecting spatial heterogeneity, measurement sensitivity, and normalization. While MERFISH data can be sparse at the single-cell level, meta cell aggregation leads to less sparse and more stable distributions.

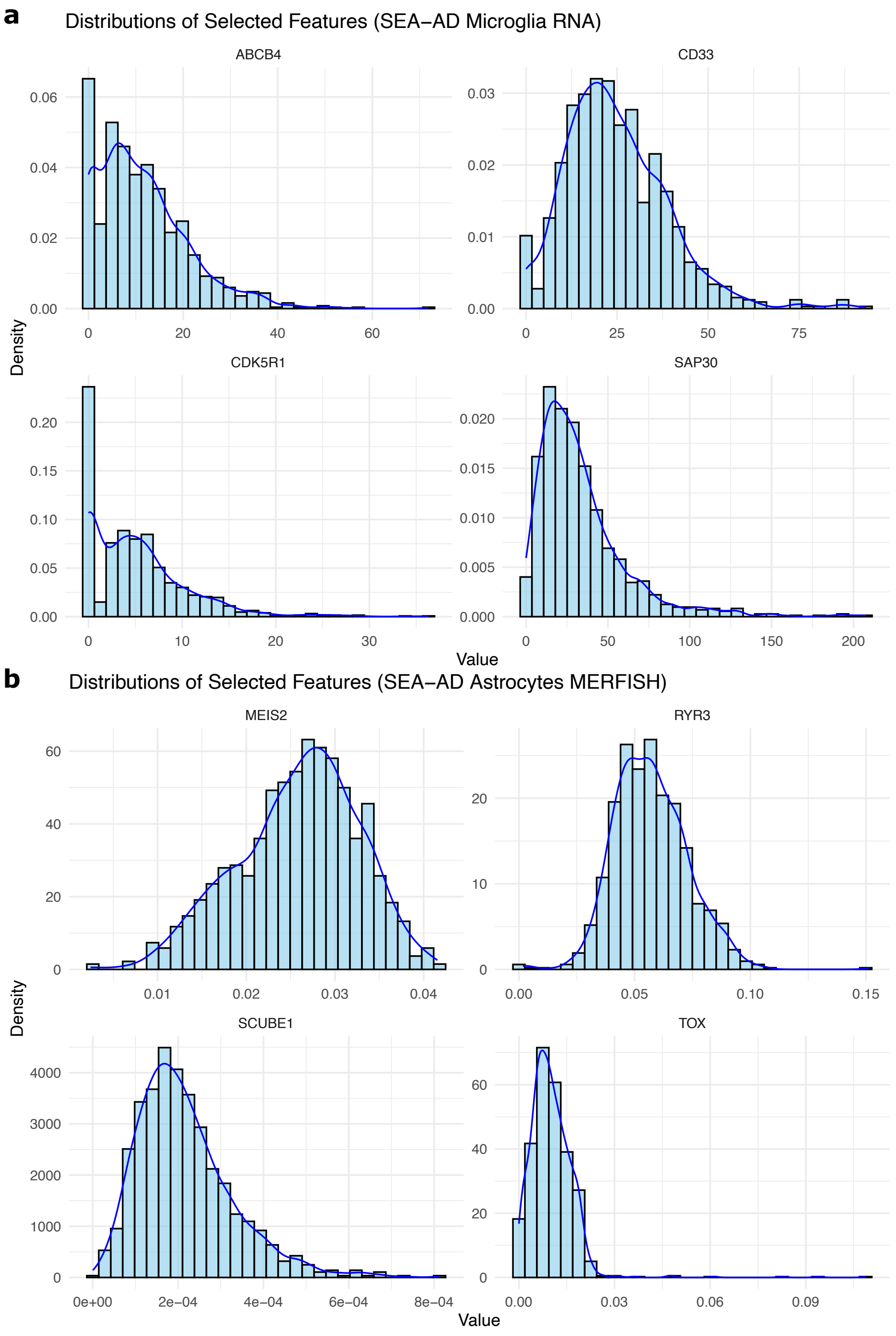


**Supplementary Fig. 7: Distributions of microglia scRNA-seq and astrocytes MERFISH data.**
**a** Distributions of four gene expressions (microglia scRNA-seq). **b** Distributions of four gene expressions (astrocytes MERFISH).

**Supplementary Table 1: Performance comparison across COSIME and other methods (Simulated high signal data).**

| Method | Binary | | | Continuous |
|---|---|---|---|---|
| | AUROC | AUPRC | Accuracy | MSE |
| Regression A | 0.818 ± 0.004 | 0.823 ± 0.002 | 0.744 ± 0.010 | 88.108 ± 1.722 |
| Regression B | 0.821 ± 0.011 | 0.828 ± 0.013 | 0.721 ± 0.013 | 72.252 ± 0.774 |
| COSIME Single-modality A | 0.827 ± 0.031 | 0.823 ± 0.044 | 0.743 ± 0.032 | 67.825 ± 4.777 |
| COSIME Single-modality B | 0.828 ± 0.024 | 0.792 ± 0.038 | 0.746 ± 0.069 | 66.658 ± 4.771 |
| **COSIME (Multi-modality Early)** | **0.845 ± 0.026** | **0.854 ± 0.029** | **0.754 ± 0.054** | **64.490 ± 4.130** |
| **COSIME (Multi-modality Late)** | **0.828 ± 0.021** | **0.853 ± 0.021** | **0.761 ± 0.019** | **54.701 ± 2.391** |
| Cooperative learning | 0.812 ± 0.038 | 0.798 ± 0.029 | 0.742 ± 0.025 | 85.798 ± 0.770 |
| DIABLO | 0.779 ± 0.049 | 0.779 ± 0.058 | 0.696 ± 0.045 | 106.943 ± 1.432 |
| MOFA+ + logistic regression | 0.719 ± 0.098 | 0.716 ± 0.065 | 0.653 ± 0.071 | 78.817 ± 10.567 |
| Aligned OT + logistic regression | 0.775 ± 0.097 | 0.742 ± 0.088 | 0.690 ± 0.121 | 67.692 ± 4.856 |
| SCOT OT + logistic regression | 0.815 ± 0.010 | 0.841 ± 0.023 | 0.737 ± 0.022 | 66.137 ± 3.345 |

**Supplementary Table 2: Performance comparison across COSIME and other methods (Transcriptomics and metabolomics from ROSMAP)**

| Method | AUROC | AUPRC | Accuracy |
|---|---|---|---|
| Logistic regression (Transcriptomics) | 0.701 ± 0.001 | 0.726 ± 0.002 | 0.654 ± 0.009 |
| Logistic regression (Metabolomics) | 0.679 ± 0.011 | 0.699 ± 0.006 | 0.594 ± 0.070 |
| COSIME Single-modality A | 0.771 ± 0.012 | 0.813 ± 0.129 | 0.728 ± 0.027 |
| COSIME Single-modality B | 0.744 ± 0.003 | 0.744 ± 0.003 | 0.786 ± 0.007 |
| **COSIME (Multi-modality Early)** | **0.792 ± 0.021** | **0.810 ± 0.022** | **0.732 ± 0.025** |
| **COSIME (Multi-modality Late)** | **0.807 ± 0.013** | **0.832 ± 0.016** | **0.723 ± 0.027** |
| Cooperative learning | 0.711 ± 0.019 | 0.730 ± 0.021 | 0.654 ± 0.017 |
| DIABLO | 0.672 ± 0.003 | 0.709 ± 0.006 | 0.610 ± 0.016 |
| MOFA+ + logistic regression | 0.722 ± 0.013 | 0.764 ± 0.013 | 0.654 ± 0.024 |
| Aligned OT + logistic regression | 0.789 ± 0.033 | 0.787 ± 0.039 | 0.719 ± 0.032 |
| SCOT OT + logistic regression | 0.783 ± 0.017 | 0.786 ± 0.038 | 0.713 ± 0.022 |

**Supplementary Table 3: Run time (in minutes) of COSIME on different datasets and systems**

| # of samples | # of features | Apple M1 Max | Intel Xeon Gold 6140 system (36 cores, 200 GB RAM) |
|---|---|---|---|
| **1000** | **200** | **4.2** | **3.0** |
| 5000 | 200 | 12.6 | 9.0 |
| 5000 | 500 | 14.4 | 9.6 |
| 10000 | 1000 | 22.2 | 19.8 |

#### Supplementary Data 1

Table showing the prediction performance of COSIME models and benchmark models for binary outcome classification.

#### Supplementary Data 2

Table showing the prediction performance of COSIME models and benchmark models for continuous outcome classification.

#### Supplementary Data 3

Feature importance and interactions for the simulated binary high signal early fusion.

#### Supplementary Data 4

Feature importance and interactions for simulated binary low signal early fusion.

#### Supplementary Data 5

Feature importance and interactions for the simulated continuous high signal early fusion.

#### Supplementary Data 6

Feature importance and interactions for the simulated continuous low signal early fusion.

#### Supplementary Data 7

Feature importance and interactions for the ROSMAP binary classification early fusion.

#### Supplementary Data 8

Feature importance and interactions for the SEA-AD scRNA-seq and scATAC-seq (oligodendrocytes) early fusion.

#### Supplementary Data 9

Feature importance and interactions for the SEA-AD scRNA-seq (microglia) and MERFISH (astrocytes) early fusion.

**Algorithm 1:** COSIME Model Training

**Training Loop**

**for** $m = 1$ **to** $M$ **do**

**Input:**

- Training: $X_1 \in \mathbb{R}^{n \times px_1}$, $X_2 \in \mathbb{R}^{n \times px_2}$
- Validation: $(X_1^{\text{val}}, X_2^{\text{val}}, y_1^{\text{val}}, y_2^{\text{val}})$
- Test: $(X_1^{\text{test}}, X_2^{\text{test}}, y_1^{\text{test}}, y_2^{\text{test}})$

**Initialization:**

- Set training iterations $M$, optimizer, and early stopping criteria
- Set tuning hyperparameters:
  batch_size, learning_rate, learning_gamma, $\lambda_{\text{KLD\_A\_weight}}$, $\lambda_{\text{KLD\_B\_weight}}$, $\lambda_{\text{OT\_weight}}$, $\lambda_{\text{CL\_weight}}$, dropout, dim, $p_{\text{earlystop\_patience}}$, $\delta$, decay

**Forward Pass:**

- Compute KLD Losses, LOT Loss, and Prediction Loss: $\mathcal{L}_{\text{KLD}_A}(\theta)$, $\mathcal{L}_{\text{KLD}_B}(\theta)$, $\mathcal{L}_{\text{LOT}}(\theta)$, $\mathcal{L}_{\text{pred}}(\theta)$
- Compute Total Weighted Loss: $\mathcal{L}_{\text{Total\_weighted}}(\theta)$

**Backpropagation:**

- Backpropagate gradients for all parameters, including the transport plan for $\mathcal{L}_{\text{LOT}}$
- Perform optimization step: $\theta \leftarrow \theta - \eta \nabla_\theta \mathcal{L}_{\text{Total\_weighted}}(\theta)$

**Early Stopping:**

Compute validation prediction loss: $\mathcal{L}_{\text{pred}}(\theta)$ for validation

**If** $\mathcal{L}_{\text{pred}}(\theta) < \mathcal{L}_{\text{pred}}(\theta_{\text{best}})$ **(improvement in validation loss)**:

`Update` $\theta_{\text{best}} = \theta$

`Reset patience counter:` $p_{\text{counter}} = 0$

**Else If** $p_{\text{counter}} > p_{\text{earlystop\_patience}}$ **(early stopping)**:

`Break training loop`

**Else**:

`Increment patience counter:` $p_{\text{counter}} \leftarrow p_{\text{counter}} + 1$

**End If**

**end for**

**Holdout Evaluation**

Evaluate best model on the test set: compute losses and metrics;

**Output**

Trained best COSIME model and the evaluation metrics;

**Algorithm 2:** COSIME Feature Importance and Interaction Computation

**Input**

$X \in \mathbb{R}^{N \times F}$: Input feature matrix,
$y \in \mathbb{R}^{N}$: Target labels

**Initialization**

- Define the model $f$ and input matrix $X$ with $N$ samples and $F$ features.
- Set number of Monte Carlo iterations $M$.
- Define the batch size, or compute it based on available memory.
- Compute the number of batches: $B = \lceil \frac{N}{\text{batch size}} \rceil$.
- Compute the memory required for each batch:

$$\text{Batch memory (GB)} = \frac{\text{batch size} \times \text{element size of } X \times F}{1 \times 10^9}.$$

- If batch memory exceeds available memory, adjust the batch size dynamically.

**Feature Importance Computation**

1. For each feature $i$ in $X$, compute the marginal contribution over $M$ Monte Carlo iterations.
2. Store feature importance values in matrix $\mathcal{S} \in \mathbb{R}^{N \times F}$:
   $\mathcal{S}_{ij} = \phi_j(s_i)$, where $\phi_j(s_i)$ is the feature importance value for feature $j$ for sample $s_i$.

**Feature Interaction Computation**

1. For each pair of features $(i, j)$, compute the interaction effect over $M$ Monte Carlo iterations.
2. Store interaction effects in matrix $\mathcal{I} \in \mathbb{R}^{F \times F}$:
   $\mathcal{I}_{ij} = \mathcal{I}_{ji}$, where $\mathcal{I}_{ij}$ is the interaction effect between features $i$ and $j$.

**Output**

$\mathcal{S} \in \mathbb{R}^{N \times F}$: Feature importance matrix,
$\mathcal{I} \in \mathbb{R}^{F \times F}$: Feature interaction matrix.
